# Species abundance distributions should underpin ordinal cover-abundance transformations

**DOI:** 10.1101/535948

**Keywords:** aggregated, beta regression, Braun-Blanquet, growth form, midpoint, ordinal transformation, species abundance distribution, sPlot, summed foliage cover, VegBank, vegetation cover

## Abstract

The cover and abundance of individual plant species have been recorded on ordinal scales for millions of plots world-wide. Many ecological questions can be addressed using these data. However ordinal cover data may need to be transformed to a quantitative form (0 to 100%), especially when scrutinising summed cover of multiple species. Traditional approaches to transforming ordinal data often assume that data are symmetrically distributed. However, skewed abundance patterns are ubiquitous in plant community ecology. A failure to account for this skew will bias plant cover estimates, especially when cover of multiple species are summed. The questions this paper addresses are (i) how can we estimate transformation values for ordinal data that accounts for the underlying right-skewed distribution of plant cover; (ii) do different plant groups require different transformations and (iii) how do our transformations compare to other commonly used transformations within the context of exploring the aggregate properties of vegetation? Using a continuous cover dataset, each occurrence record was mapped to its commensurate ordinal value, in this case, the ubiquitous Braun-Blanquet cover-abundance (BBCA) scale. We fitted a Bayesian hierarchical beta regression to estimate the predicted mean (PM) cover of each of six plant growth forms within different ordinal classes. We illustrate our method using a case study of 2 809 plots containing 95 812 occurrence records with visual estimates of cover for 3 967 species. We compare the model derived estimates to other commonly used transformations. Our model found that PM estimates differed by growth form and that previous methods overestimated cover, especially of smaller growth forms such as forbs and grasses. Our approach reduced the cumulative compounding of errors when transformed cover data were used to explore the aggregate properties of vegetation and was robust when validated against an independent dataset. By accounting for the right-skewed distribution of cover data, our alternate approach for estimating transformation values can be extended to other ordinal scales. A more robust approach to transforming floristic data and aggregating cover estimates can strengthen ecological analyses to support biodiversity conservation and management.

## Introduction

Field-based assessment of the cover and abundance of individual plant species is complex. Observers making on-ground visual estimates of plant cover need to account for, and assess, foliage cover of different densities, dimensions, shapes and structures across multiple species, growth forms and strata. So too, counting cryptic, clonal, or copious numbers of plants can be complicated. Owing to this complexity, vast numbers of floristic plots across many continents have been surveyed using ordinal scales (Schaminée et al. 2009; Dengler et al. 2011; Chytrý et al. 2016). Whilst, in Braun-Blanquet (1932) originally described an abundance-dominance scale, the practical, on-ground application of this scale is to assess plant cover, and where cover is less than 5%, abundance is also assessed. The Braun-Blanquet cover-abundance (BBCA) scale is perhaps the most common ordinal scale used in plant ecology. For example, within the vegetation plot database sPlot v2.1 (www.idiv.de/splot), more than 745 000 plots (66%) have recorded plant occurrence using Braun-Blanquet cover-abundance (sPlot extract supplied by Borja Jiménez-Alfaro, 19^th^ September 2017). This volume of data is testament that ground-based visual assessments of cover-abundance using ordinal scales provide a cost-effective, rapid and non-destructive approach to gathering the data needed to summarise the composition and structure of plant communities. These data represent a wealth of investment in field effort and have supported major advances in vegetation classification, mapping and distribution modelling.

The ever-growing access to global vegetation plot databases (Dengler et al. 2011; Schaminée et al. 2011) has opened pan-continental opportunities to explore many uses of floristic data. Some ecological questions may best be addressed using aggregate properties of vegetation, such as the summed total foliage cover within a plot or across strata, the total summed cover or abundance of exotic or invasive species, or the relative cover or abundance of plants within different functional, taxonomic or growth form groups. Summing cover to derive aggregate properties of floristic data have a multitude of uses in ecology including assessing presence and diversity of faunal habitat, as covariates in species’ distribution models (SDMs), for assessing the spatial and temporal status of ecosystem baselines, predicting the effects of shifts in climate, land use and land cover, or measuring site-scaled responses to disturbance (e.g. Scholes & Biggs 2005; McElhinny et al. 2006; Pereira et al. 2010). Aggregate properties of vegetation data are particularly relevant to exploring ecological questions concerning the patterns, processes and prognoses at a range of spatial scales in contemporary and predicted future landscapes.

There are many applications where ordinal data have been used successfully, such as ordination, classification, modelling or mapping of vegetation communities (e.g. Podani 2005; Podani 2006; Lyons et al. 2016) and for modelling the cover of single species (e.g. Damgaard 2014; Irvine et al. 2016). However, ordinal scaled cover observations of individual species cannot be summed (Guisan & Harrell 2000; Podani 2006; Chen et al. 2008b) and need to be transformed into a continuous scale prior to aggregating.

Approaches to transforming Braun-Blanquet cover-abundance (BBCA) ordinal data have been proposed by Tüxen and Ellenberg (1937) and Braun-Blanquet (1964) (see Table 3 in van der Maarel 1979). In addition, van der Maarel (1979) proposed the ordinal transform value (OTV) with different scale adjustments, as a solution for converting ordinal data to percentage cover values. All these methods tend to transform data to the approximate midpoint of the ordinal class range for observations of cover greater than 5%. For classes with cover less than 5%, the transformation values appear arbitrary and differ considerably (Table 1 columns 4–6).

**Table 1:** Class divisions for the 1–6 Braun-Blanquet ordinal cover-abundance (BBCA) scale (columns 1–3), previous proposals for transforming them to percentage cover (columns 4–6), and proposed transforms (independent of growth form) based on estimating the predicted mean (PM) from a beta distribution of observed quantitative cover data and the lower 2.5% and upper 97.5% credible intervals. Number of observations (n) for BBCA1 (n = 54 811); BBCA2 (n = 26 968); BBCA3 (n = 11 946); BBCA4 (n = 1583); BBCA5 (n = 366) and BBCA6 (n = 138).

| Column 1 | 2 | 3 | 4 | 5 | 6 | 7 | 8 |
| --- | --- | --- | --- | --- | --- | --- | --- |
| BBCA Class | Range of cover (%) | Qualitative abundance terms | Tüxen & Ellenberg (1937) <sup>1</sup> | Braun-Blanquet (1964) <sup>1</sup> | van der Maarel (2007) <sup>2</sup> | PM | Credible interval |
| 1 | <5 | e.g. present, few, rare, erratic, occasional, uncommon | 0.1 | 0.1 | 1 | 0.49 | 0.48–0.51 |
| 2 | <5 | e.g. common, abundant, many, several | 2.5 | 5 | 2 | 0.74 | 0.72–0.76 |
| 3 | 5–25 |  | 15 | 17.5 | 17.5 | 8.95 | 8.84–9.07 |
| 4 | 26–50 |  | 37.5 | 37.5 | 35 | 38.77 | 37.97–39.57 |
| 5 | 51–75 |  | 62.5 | 62.5 | 70 | 62.43 | 60.69–64.13 |
| 6 | 76–100 |  | 87.5 | 87.5 | 140 | 81.24 | 79.10–83.26 |

<sup>1</sup> adapted from van der Maarel (1979).
<sup>2</sup> ordinal transform values (OTV) using 1.415 weighting factor (van der Maarel 2007).
Column 3 shows some of the qualitative descriptors used by field surveyors to divide observations between BBCA1 and BBCA2.

Transforming data to the approximate midpoint of the class ranges assumes that data are symmetrically distributed within each class. Yet, patterns in plant abundance including density, biomass (Chiarucci et al. 1999; Morlon et al. 2009), frequency (Chiarucci et al. 1999), percentage cover (Damgaard 2009), size, energy use and productivity (Whittaker 1965) have all been shown to have a right-skewed distribution; skewed species abundance distributions occur in every known multi-species community (McGill et al. 2007). Midpoint transformations are inflexible to the underlying distribution of cover data and assume that the distribution does not vary across species, groups of plant entities (such as growth forms, life forms, functional or taxonomic groups), vegetation types or biomes. Due to the prevalence of right-skewed distribution, we predict that midpoint transformations overestimate cover and the compounding of these errors will result in gross overestimation of summed cover for aggregated properties.

Here we develop a flexible approach to estimate cover transformations for ordinal scaled data that can then be used to provide accurate estimates of summed vegetation cover. The method we describe is applicable to data in any ordinal scale, can be extended to allow for differences in vegetation type or among biomes and can accommodate alternative aggregate properties of plant data such as growth forms, life forms, functional or taxonomic groups. To demonstrate the potential applicability of our approach we build and then validate the model using two separate and independent datasets.

Given that diverse architectures and spatial arrangements of foliage lead to varied patterns of plant cover (Damgaard 2013), we also predict that different plant growth forms will require different transformation values. Growth forms are practical and observable entities that can inform site-based assessment and monitoring, are recognizable from remotely-sensed imagery and are used to report on broad-scale biodiversity assessment or baselines (Pereira et al. 2013) with which we can measure change in cover (Pettorelli et al. 2014; Abelleira Martínez et al. 2016).

## Materials and Methods

We outline the key steps required to estimate transformation values within ordinal classes for different plant groups. A pre-requisite for our method is cover data that have been collected on a continuous cover scale, ideally sourced from the same study region and vegetation types as the ordinal cover data. To prepare the input data for the model, ordinal values need to be mapped, *a posteriori*, to this continuous cover data as an intermediary variable (Figure 1, Step 1). Models, with a beta distribution, are then used to predict the mean cover of each plant group within each ordinal cover class. This predicted mean cover is the transformation value (Figure 1, Step 2). Using a case study, we explore summed cover estimates for different plant groups and evaluate the performance of the ordinal cover transformations. We compare our transformation to existing approaches in the context of summed cover for plant groups (Figure 1, Step 3). We evaluate the robustness of our predicted mean transformations on an independent dataset (Figure 1, Step 4).

**Figure 1:**
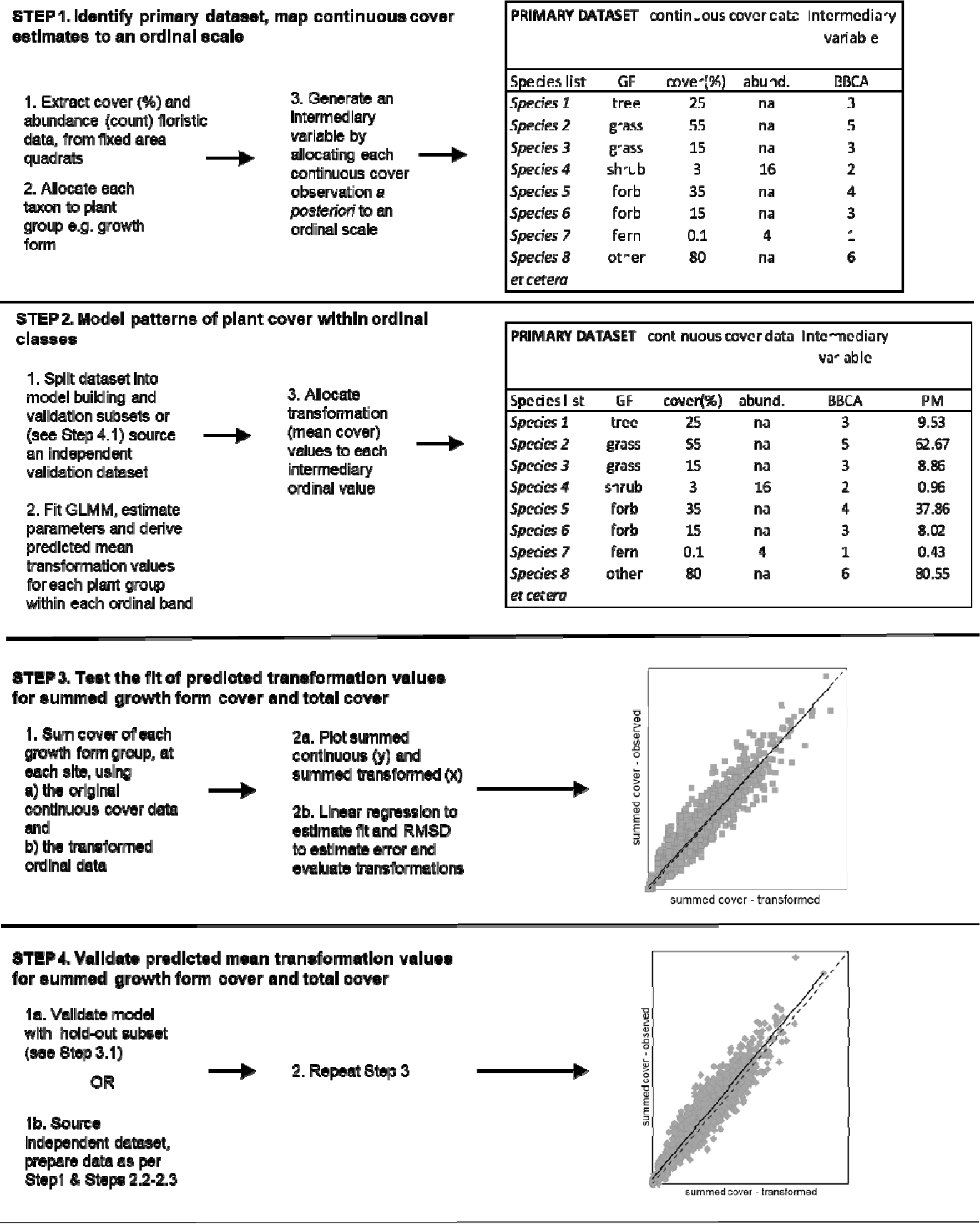
Workflow showing the major elements required to estimate transformation values for ordinal data using continuous cover estimates. Here we use Braun-Blanquet cover-abundance (BBCA) 1-6 scale, although this approach could be extended to any ordinal scale. Note, this flow diagram represents data from one plot, but many plots are needed to obtain robust estimates of mean cover.

### Estimate mean cover using parameters of a beta distribution

We used a generalised linear mixed model (GLMM) with a beta distribution to derive estimates of the mean vegetation cover, within an ordinal class, given a plant’s growth form and random variation owing to plot identity. Individual species cover are continuous proportional estimates, and once suitably transformed, fall within the known range (0<y<1). Linear regression with a normal distribution is inappropriate for the analysis of proportions, such as percent plant cover, because data often violate assumptions such as normality and homogeneity of errors and furthermore fitted values can fall outside of the range [0,1] (Ferrari & Cribari-Neto 2004). A common approach to address these problems is to apply arcsine or logit transformations to the response variable, prior to regression (Warton & Hui 2011), although the results can be difficult to interpret (Ferrari & Cribari-Neto 2004).

Numerous authors have instead demonstrated that percent plant cover are more appropriately analysed by assuming that cover approximates a two-parameter beta distribution (Ferrari & Cribari-Neto 2004; Chen et al. 2008a; Cribari-Neto & Zeileis 2010; Herpigny & Gosselin 2015). Beta distributions are attractive because fitted values are constrained between the interval 0<y<1 and they can accommodate asymmetrical distributions with left- or right-skew. This flexibility makes beta distributions highly suitable for modelling diverse and often asymmetrical plant cover data (Cribari-Neto & Zeileis 2010).

We present a Bayesian GLMM with a logit link to estimate the parameters of the beta distribution and allowed these parameters to vary among ordinal classes and plant growth forms. Estimates of these parameters were used to derive the predicted mean (PM) for each plant growth form in each ordinal class.

The proportional vegetation cover is given by the two-parameter beta distribution;

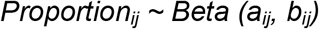

Where *a_ij_* and *b_ij_* are shape parameters for species *j* in plot *i, and i = 1,…n plots*. The shape parameters are further defined as

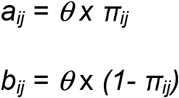

where *θ* allows for potential overdispersion to be incorporated in the model (Zuur et al. 2013).

π_*ij*_ is modelled with a logit link

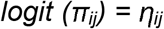

The model consists of regression parameters (β) for each ordinal class, plant growth form and their interactions, plot level random intercepts and variance (*Zi*):

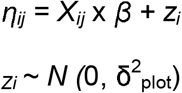

Where *Z_i_* is a random intercept for plot, *X_ij_* are the matrix of all covariates (ordinal classes and their interaction with plant growth form) and *β* are the regression parameters for each covariate. That is, for each ordinal class 1…6, separate *β* were estimated for each plant growth form. For a simplified example with two growth forms and two ordinal classes this can also be expressed as:

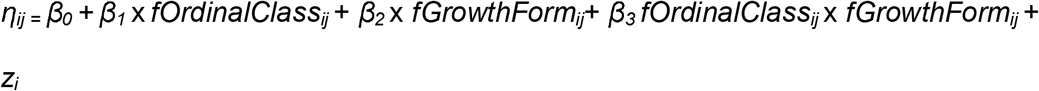

Where β_0_ = predicted value of logit transformed cover if species *j* belongs to the “reference” growth form and its’ value in plot *i* has the “reference” level ordinal cover-abundance class.

β_1_ = departure of the predicted value for species *j* from β_0_ if the observation is of another ordinal cover-abundance class.

β_2_ = departure of predicted value from β_0_ if species *j* belongs to another growth form.

β_3_ = departure of predicted value from β_0_ + β_1_+ β_2_ when neither growth form nor ordinal cover-abundance class are of the reference level.

In this example, *fOrdinalClass_ij_* and *fGrowthForm*_ij_ are binary dummy variables coding growth form and cover-abundance scale categories, thus *Xij* is a vector containing values for these dummy variables (including their products) for species *j* in plot *i*.

We included plot as a random intercept because although we assumed each plot should follow the characteristic skewed species abundance curve, we expected variation among plots and hence differences in the average cover of any given ordinal class and plant growth form.

This basic model structure can be easily expanded to accommodate other possible sources of variation, such as among vegetation types or owing to the richness of plant species within a plot. In this case study, we decided not to include additional covariates to minimise computational demands and simplify model interpretation and operational complexity.

The model was fit via Markov chain Monte Carlo optimization in JAGS (http://mcmc-jags.sourceforge.net) via the R2jags package (Su & Yajima 2015) within R 3.5.0 (R Core Team 2018). Posterior parameter estimates and back-transformed predicted means were derived from 3 chains, with a burn-in of 3000 iterations, 15 000 subsequent iterations per chain and with a thinning rate of 15. Autocorrelation and mixing were visually inspected. The interaction models were compared to additive models using Deviance Information Criteria. Appendix S1 contains R code for our models.

### Case study – New South Wales, Australia

We illustrate our model with a case study where we have used 1-6 BBCA as our ordinal scale and grouped plants into six growth form categories. Following is a brief description of how we prepared the case study dataset to build our model. We note that randomly generated data from an appropriate beta distribution (for similar example see Damgaard 2014) could also be used to demonstrate our approach. However, we chose to use a large archival dataset from a range of bioclimatic regions and vegetation types to demonstrate that, despite the underlying variation, our approach still led to robust estimates of summed cover.

#### 1. Preparation of observed percentage cover dataset

To demonstrate our modelled approach, we sourced case study data from archival quantitative floristic data that met three considerations: (i) each species record included a visual estimate of foliage cover on a continuous scale from 0.1% to 100% and a count of abundance where cover was less than 5%; (ii) in each plot, full species inventories were recorded from a fixed-area (400 m^2^) and (iii) sites covered a wide geographic distribution (Appendix S2—Figure 1) and included a wide range of vegetation types with different structural complexity including rainforests, forests, woodlands, shrublands, grasslands and wetlands (Keith 2004). A total of 2809 geo-referenced plots containing 95 812 occurrence records with visual estimates of cover for 3967 species met these criteria and were exported from the NSW BioNet Atlas database (www.bionet.nsw.gov.au).

### Analysis of the empirical cover distribution

To confirm our assumption of right-skewed distribution of cover data we plotted our data and used the ‘skewness’ function in the e1071 package (Meyer et al. 2017) within R 3.3.3 (R Core Team 2018) to calculate the adjusted Fisher-Pearson skewness coefficient (*G_1_*) (Joanes & Gill 1998) for the whole distribution, and for distributions within each BBCA class. Skewness is a diagnostic tool usually used to test the symmetry of the data distribution. Here, we interpret skewness coefficients as being strongly and positively skewed when the *G_1_* coefficient is greater than 0.5 (Bulmer 1979; Doane & Seward 2011).

#### 2. Preparation of plant group entities

All taxa were allocated to one of six growth form categories: tree, shrub, grass and grass-like (hereafter referred to as grass), forb, fern and other (remaining growth forms) (Oliver et al. submitted). For each growth form in each plot, total cover was estimated by summing the observed quantitative estimates of cover and the estimates of cover derived from the transformations of the ordinal data.

#### 3. Allocating an intermediary variable

We created an intermediary variable by matching each quantitative estimate of cover for every floristic record (n = 95 812) to its commensurate ordinal value. Any ordinal scale can be used to partition data, but here we demonstrate our approach by allocating data to 1–6 BBCA (Table 1). BBCA1 and BBCA2 were assigned based on their observed foliage cover (<5%) and abundance; where BBCA1 ≤ 10 and BBCA2 > 10 individuals. The pragmatic choice of ten individuals provides an explicit quantitative abundance threshold between classes BBCA1 and BBCA2. BBCA3–BBCA6 were assigned based on observed foliage cover (≥5%) (Mueller-Dombois & Ellenberg 1974). The ordinal dataset created by this process approximates the form of many data held within vegetation databases.

In our case study dataset, observations of 5% cover were more prevalent than expected from a typical theoretical beta distribution (Figure 2). This bias was detected in preliminary model convergence diagnostics and model fit suggested that, for our case study, it would be preferable to split the data and separately model (i) BBCA1 and BBCA2 bounded between 0 and less than 5% cover and (ii) BBCA3 to BBCA6 bounded between 5% and 100% cover inclusive. To ensure the response variable was bounded by 0 and 1, percent cover was transformed using (y-a)/(b-a) where in (i) a = 0 and b = 5 and in (ii) a = 5 and b = 100 (Cribari-Neto & Zeileis 2010). In the second model, the response variable was further transformed using (y * (n-1))/n where n = sample size (Cribari-Neto & Zeileis 2010). This split-model approach may not be necessary for all datasets, especially where data are derived from less subjective cover methods (e.g. point intercept or pin frame) but is included here to support the handling of datasets with similar patterns in distribution (see Appendix S3—Figures S1–S4 for other datasets that appear to show similar pattern).

**Figure 2:**
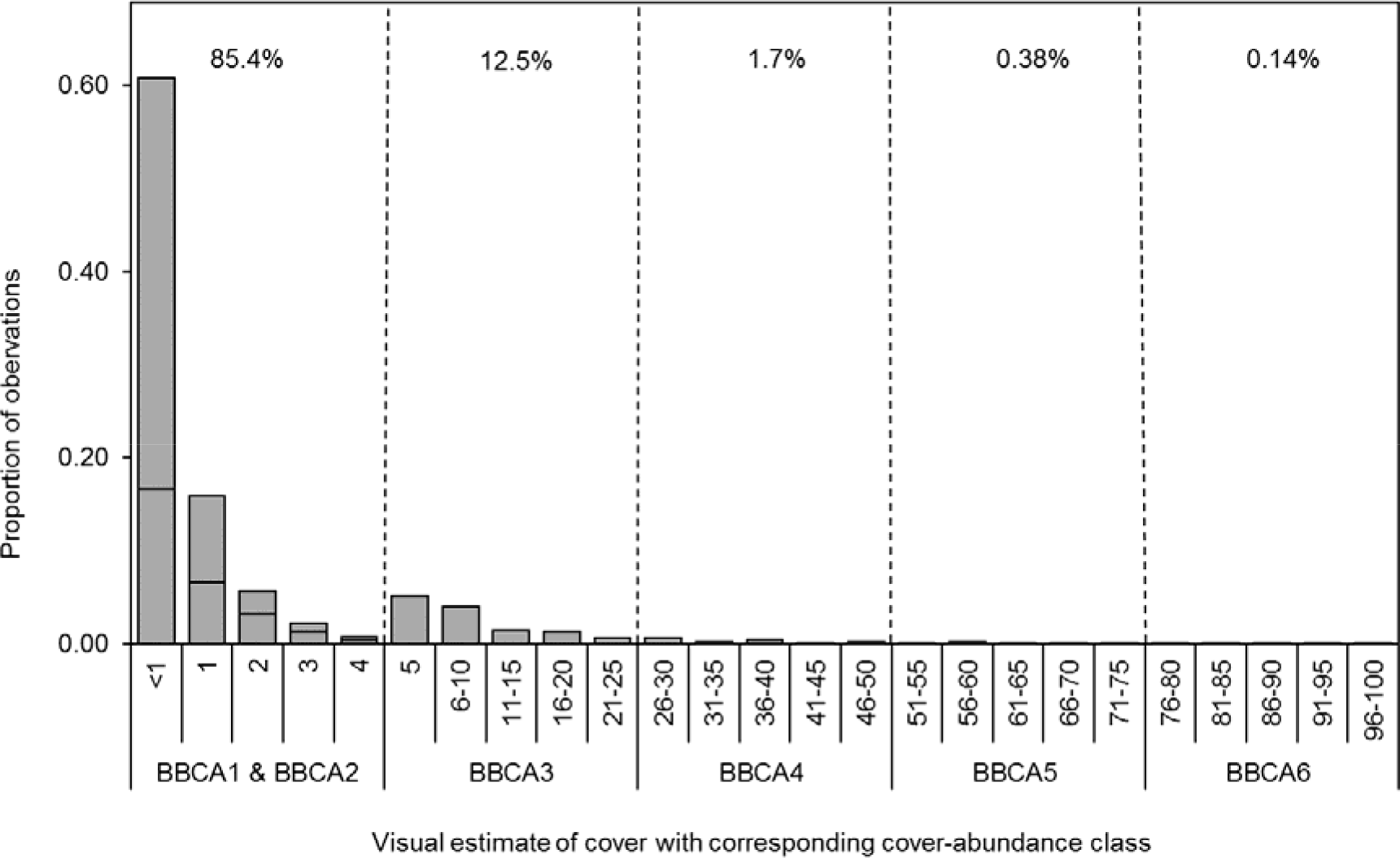
Distribution of visual estimates of cover for 95 812 observations, and their corresponding Braun-Blanquet cover-abundance (BBCA) class for our case study. Dashed vertical lines show cut points between each BBCA class. Number of observations (n) for BBCA1 (n = 54 811); BBCA2 (n = 26 968); BBCA3 (n = 11 946); BBCA4 (n = 1 583); BBCA5 (n = 366) and BBCA6 (n = 138). Numbers between the dashed lines show the percentage of each class in the dataset. BBCA1 and BBCA2 (both represent <5% cover) are shown as stacked histograms; BBCA1 (≤ 10 individuals) sits above BBCA2 (> 10 individuals). See Appendix S3—Figures 1-4 for comparison with other archival datasets.

### Evaluation of past and proposed approaches to transforming ordinal data

We transformed each of the 1–6 BBCA records using three different approaches outlined in Table 1. We then evaluated these past approaches proposed by Tüxen and Ellenberg (1937), Braun-Blanquet (1964) and van der Maarel (2007) to the PM estimated from a beta distribution.

For each plot, growth form cover and total cover were calculated by summing the observed continuous cover estimates (%) and the estimates of cover derived from the various transformations. Linear regression models with zero-intercept were fitted to the sum of observed continuous cover data (y) and sum of transformed cover data (x) cover data in R 3.5.0 (R Core Team 2018). We can justify using a regression through the origin because we are most interested in comparing the slope of the regression line to the 1:1 line of best fit to determine if our PM models were over or underpredicting summed cover. We compared the root mean squared deviation (RMSD) (see Eq. 1) as an estimate of the deviation of the transformed cover values from the 1:1 line.

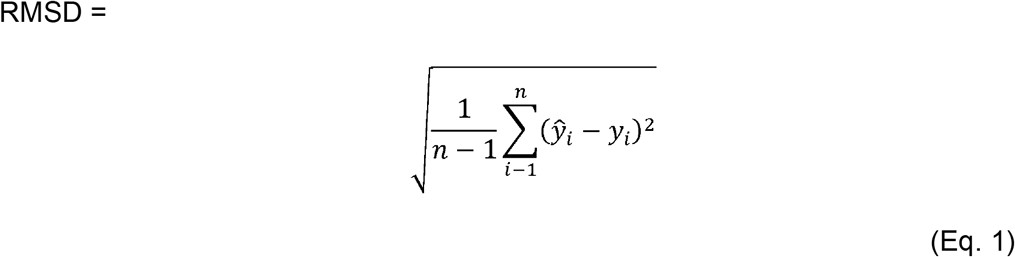

Where 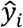 are the predicted cover values; *y*_*i*_ are the observed cover values and n is the number of observations.

The RMSD estimate represents the mean deviation of transformed cover values with respect to the true observed cover values. We also compared estimates of the slope with lower and upper 95% confidence intervals expecting that robust transformations would result in a slope = 1 and transformations that overestimate summed cover will have a slope <1. We include the adjusted coefficient of variation (R^2^) to evaluate how much of the linear variation of observed cover values is explained by the variation of transformed cover values.

We note that RMSD is useful for evaluating models as it represents an absolute measure of fit to the 1:1 line and reports the prediction error in the same units as the data (i.e. summed cover). Whereas adjusted R^2^ gives a relative measure of proportion of total variance that is explained by the model on a scale between 0 and 1.

We validated the PM transformation values on an independent dataset (2 227 sites with 51 497 observations) from West Virginia Natural Heritage Program (Vanderhorst et al. 2012) accessed from VegBank (Peet et al. 2013 accessed 28th Aug 2018). Whilst VegBank has a primary role for enabling the vegetation classification, large volumes of individual floristic observations are available for ecoinformatic synthesis and analysis. Owing to the ease of access and completeness of datasets stored in VegBank we were able to validate our model estimates on a geographically distinct dataset containing cover estimates of plants from entirely different vegetation communities. Details outlining the data preparation are included in Appendix S6.

## Results

### The empirical cover distribution

The source continuous cover data were right-skewed and dominated by low cover—85% of observations were between 0.1 and 4%, and 60% of these observations were of cover less than 1% (Figure 2). Data were heavily right-skewed for the whole distribution (*G_1_* = 5.62) and right-skewed within five of the six BBCA classes (BBCA1 *G_1_* = 2.64, BBCA2 *G_1_* = 1.57, BBCA3 *G_1_* = 1.04, BBCA4 *G_1_* = 0.61 and BBCA6 *G_1_* = 0.95). Only BBCA5 had a skewness coefficient less than 0.5 (*G_1_* = 0.36). We also note potential observer bias for 5% cover. These patterns are similar to other visually estimated floristic cover data from other archived datasets (see Appendix S3—Figures 1-4).

### Estimate mean cover using parameters of a beta distribution

Table 1 (columns 7 and 8) shows the predicted mean transformations and their lower 2.5% and upper 97.5% credible interval for each ordinal class, independent of growth forms. The most marked differences are noted in BBCA2 and BBCA3, where the predicted means are well below the previous approaches. The predicted mean for class BBCA6 is lower than the midpoint but was derived from relatively few observations (n = 138).

### Evaluation of past and proposed transform values for summed growth form cover

Estimates of the PM suggest that accounting for growth form within each ordinal class results in more robust summed cover estimates. Credible intervals suggest that in classes BBCA1 to BBCA3, trees typically have higher mean cover and warrant higher transformation values (Table 2). Credible intervals also suggest the need for separate transformation values for shrubs in BBCA1 and BBCA2 and a lower value for forbs in BBCA2 and BBCA3 (Table 2).

**Table 2:**
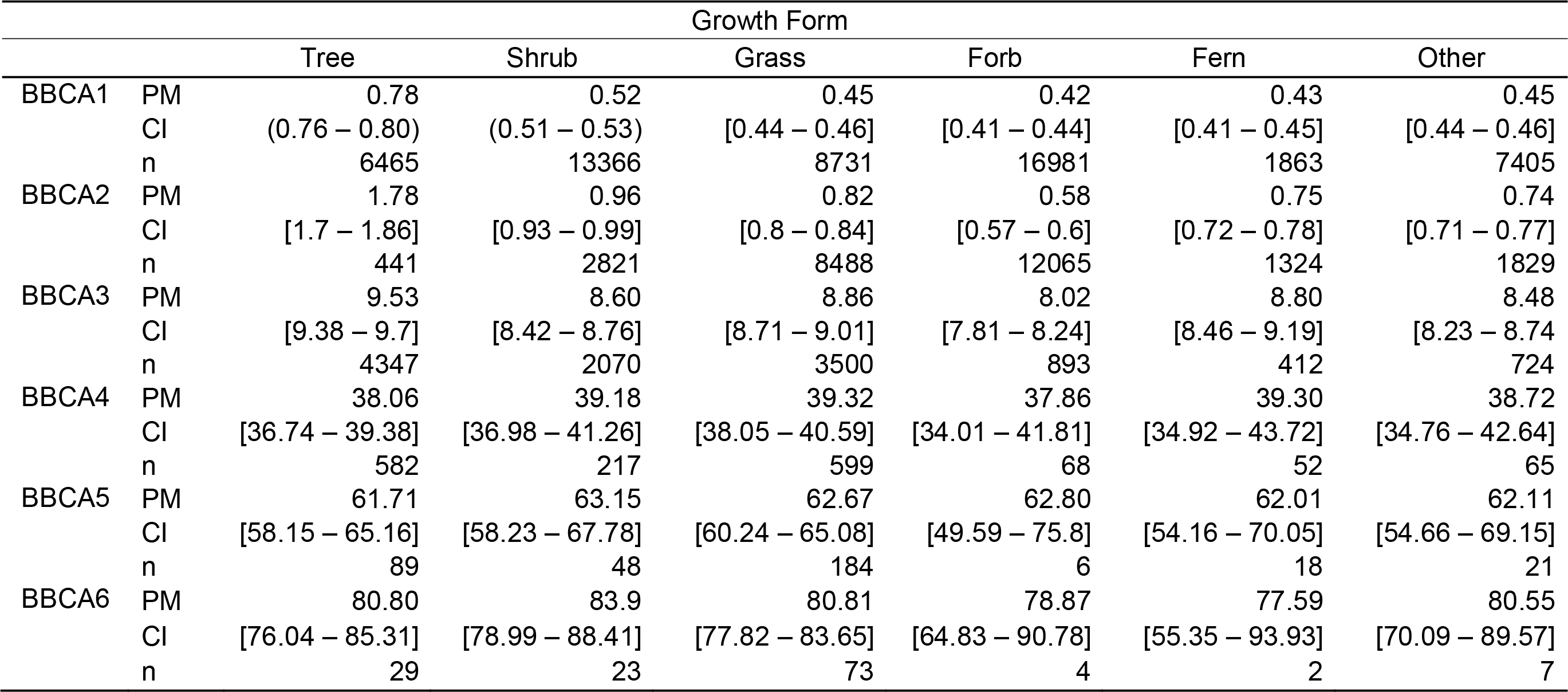
Proposed transformation values, tailored to different growth forms, based on estimates of the predicted mean (PM) from a beta distribution of observed data. Lower 2.5% and upper 97.5% credible intervals (CI) are shown in square brackets; n = number of individual observations for each Braun-Blanquet cover-abundance (BBCA) class.

|  |  | Growth Form |  |  |  |  |  |
| --- | --- | --- | --- | --- | --- | --- | --- |
|  |  | Tree | Shrub | Grass | Forb | Fern | Other |
| BBCA1 | PM | 0.78 | 0.52 | 0.45 | 0.42 | 0.43 | 0.45 |
|  | CI | (0.76 – 0.80) | (0.51 – 0.53) | [0.44 – 0.46] | [0.41 – 0.44] | [0.41 – 0.45] | [0.44 – 0.46] |
|  | n | 6465 | 13366 | 8731 | 16981 | 1863 | 7405 |
| BBCA2 | PM | 1.78 | 0.96 | 0.82 | 0.58 | 0.75 | 0.74 |
|  | CI | [1.7 – 1.86] | [0.93 – 0.99] | [0.8 – 0.84] | [0.57 – 0.6] | [0.72 – 0.78] | [0.71 – 0.77] |
|  | n | 441 | 2821 | 8488 | 12065 | 1324 | 1829 |
| BBCA3 | PM | 9.53 | 8.60 | 8.86 | 8.02 | 8.80 | 8.48 |
|  | CI | [9.38 – 9.7] | [8.42 – 8.76] | [8.71 – 9.01] | [7.81 – 8.24] | [8.46 – 9.19] | [8.23 – 8.74] |
|  | n | 4347 | 2070 | 3500 | 893 | 412 | 724 |
| BBCA4 | PM | 38.06 | 39.18 | 39.32 | 37.86 | 39.30 | 38.72 |
|  | CI | [36.74 – 39.38] | [36.98 – 41.26] | [38.05 – 40.59] | [34.01 – 41.81] | [34.92 – 43.72] | [34.76 – 42.64] |
|  | n | 582 | 217 | 599 | 68 | 52 | 65 |
| BBCA5 | PM | 61.71 | 63.15 | 62.67 | 62.80 | 62.01 | 62.11 |
|  | CI | [58.15 – 65.16] | [58.23 – 67.78] | [60.24 – 65.08] | [49.59 – 75.8] | [54.16 – 70.05] | [54.66 – 69.15] |
|  | n | 89 | 48 | 184 | 6 | 18 | 21 |
| BBCA6 | PM | 80.80 | 83.9 | 80.81 | 78.87 | 77.59 | 80.55 |
|  | CI | [76.04 – 85.31] | [78.99 – 88.41] | [77.82 – 83.65] | [64.83 – 90.78] | [55.35 – 93.93] | [70.09 – 89.57] |
|  | n | 29 | 23 | 73 | 4 | 2 | 7 |

When these growth form specific transformations were evaluated using the summed cover estimates RMSD did not exceed 9.50 (trees) (Figure 3 and Appendix S4—Table 1). In contrast, estimates based on past transformations frequently resulted in RMSD exceeding 10. Slope ranged from 0.91 (forbs) to 1.05 (others), whereas past transformations slopes were <0.85, suggesting considerable overestimation of summed cover (see Appendix S4—Table 1 and Appendix S5—Figures 1-4).

**Figure 3:**
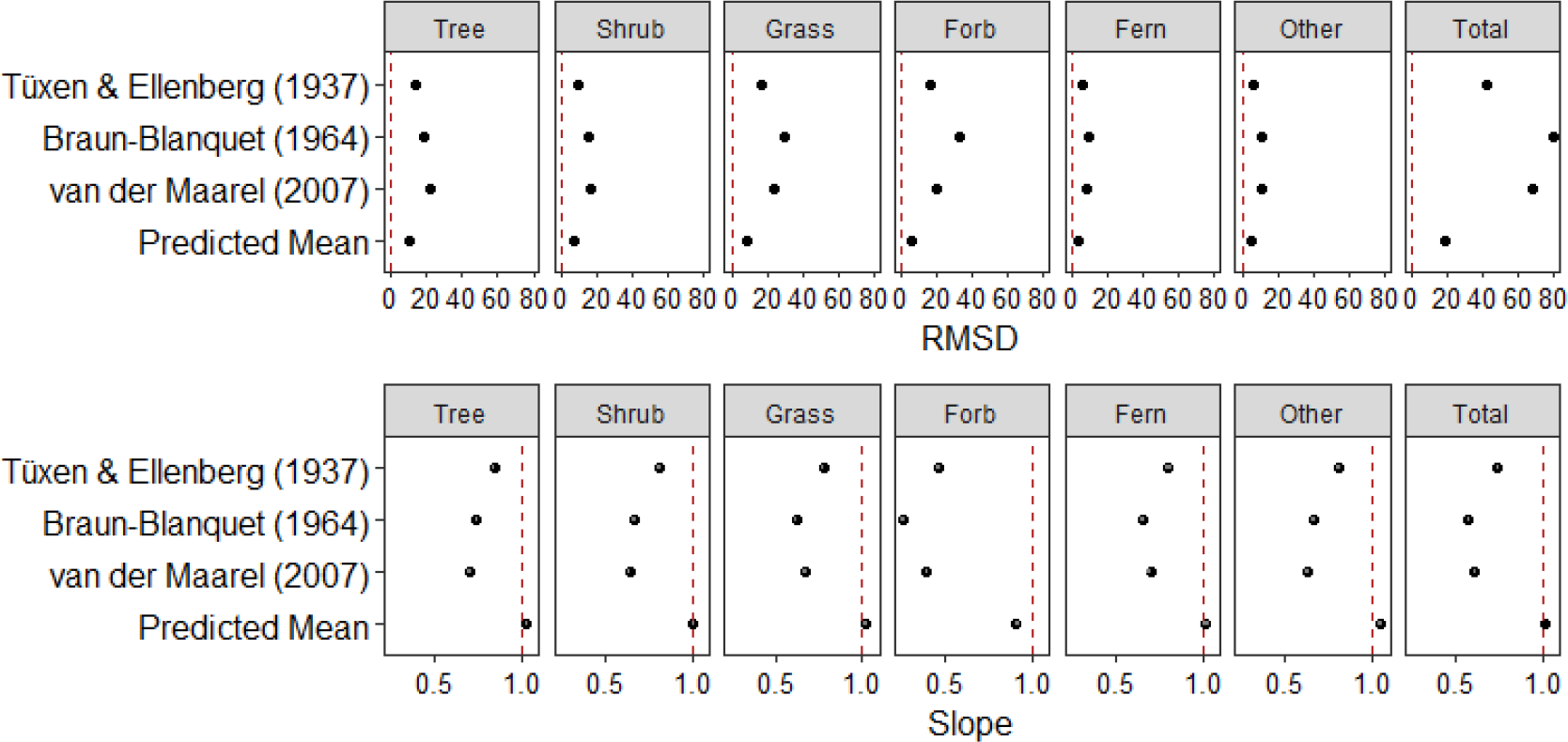
Results of linear regression with zero-intercept to compare root mean squared deviation (RMSD) and slope for each growth form and for total summed cover under previous transformations compared to the predicted mean transformation. Lower and upper 95% confidence intervals are shown for slope. Vertical dashed lines represent the perfect regression fit where RMDS = 0 and slope = 1. (data table supplied in Appendix S4—Table 1). Number of observations (n) for trees (n = 11 953); shrubs (n = 18 545); grasses (n = 21 575); forbs (n = 30 017); ferns (n = 3671); other (n = 10 051) and total (n = 2809). See Appendix S5—Figures 1a-f to 4a-f for plots of all growth forms and three previous approaches to transformation proposed by Tüxen and Ellenberg (1937); Braun-Blanquet (1964) and van der Maarel (2007).

### Evaluation of past and proposed transform values for total summed cover

Evaluation of summed total cover revealed that when transformations are tailored to growth forms, the PM performed better than existing approaches (Figure 3). The PM reduced the overestimation of total summed cover by up to 4 times. The evaluation of model fit for summed total cover using past approaches generally revealed a poorer model fit: RMSD ranged from 41.47–79.37 (PM = 18.21) (see Appendix S4—Table 1) and slope ranged from 0.57 to 0.74 (PM = 1.01) and adjusted R^2^ ranged from 0.61-0.96 (PM – 0.97).

Evaluation of the growth form specific PM transformation on an entirely independent validation dataset from West Virginia Natural Heritage Program (Vanderhorst et al. 2012) show that transformations were robust, although tended to underestimate summed cover of most growth forms (Appendix 6—Table 1). RMSD ranged between 1.59 (others) and14.97 (trees); slope ranged between 1 (others) and 1.12 (forbs) and adjusted R^2^ were high and ranged between 0.97 (trees and shrubs) and 0.93 (others). When compared to the transformation proposed by Tüxen and Ellenberg (1937), the PM transformation values were marginally better. Tuxen and Ellenberg (1937) transformation values tended to result in an overestimate summed cover of all growth forms (Appendix S7—Figures 2a-f); RMSD was consistently higher than PM transformations for all growth forms; slopes were further from 1 between 0.76 (forbs) and 0.95 (shrubs) and adjusted R^2^ ranged between 0.86 (others) and 0.98 (trees) (Appendix 6—Table 1).

Evaluation of total cover, using the PM transformation values, showed RMSD was less than that estimated if the transformation was undertaken using estimates of Tüxen and Ellenberg (20.54 *cf.* 27.01) (Appendix 6—Table 1) and PM transformation values show a slight underestimation (slope = 1.1; adjusted R^2^ = 0.98) when tested on the independent dataset.

Scatter plots showing the relationships between visual estimates of summed cover for all six growth form groups using the PM model and for Tüxen and Ellenberg (1937) transformations are provided in Appendix S7—Figures 1a-f and Figures 2a-f.

## Discussion

Transforming ordinal data to a quantitative form is common practice in plant ecology and extends across disciplines including restoration (Fill et al. 2017), classification (Cawsey et al. 2002; Faber-Langendoen et al. 2007; Wiser & De Cáceres 2013); and for assessing disturbance (Scott & Kirkpatrick 2008; Knapp & Ritchie 2016). Similarly, universal skewed patterns in the species abundance distribution are a long standing and well recognised pattern in ecology (e.g. MacArthur 1960). The data we present here are no exception. Yet the integration of these two concepts, underpinned by a robust modelling approach has received little attention, especially in the context of synthesizing information on aggregate properties of vegetation data. We demonstrate, using two large quantitative independent datasets that when the underlying right-skewed cover distribution is accounted for, a more robust set of transformations are generated. Where the aggregate properties of floristic data are of interest, our method, unlike previous approaches to transformation of ordinal data, does not overinflate cover.

Where possible, we advocate that others replicate this approach and source continuous cover data, so that the means within each ordinal class can be estimated accounting for the underlying distribution. Ideally, the continuous cover datasets will encompass the same temporal and spatial variation as that of the ordinal data. Notwithstanding these recommendations for best-practice, we have demonstrated our modelling approach can produce robust estimates of summed cover using floristic data from geographically distinct dataset containing observations of entirely unrelated vegetation communities. We expect the estimates of summed cover would further improve had we used representative data from that region and vegetation to model specific estimates of the parameters for the beta distribution. Undoubtedly there will be circumstances where appropriate continuous data will not be available and the parameters of the beta distribution cannot be estimated for a specific study or region. In these situations, adopting the PM transformations provided in Tables 1 and 2 would be preferable to application of ordinal class midpoints. When plant cover are right-skewed, midpoint transformations will bias and overestimate total cover. Hierarchical models are useful for handling complex interactions in observational data. Despite the size of the initial dataset, some plant groups were poorly represented in the higher cover classes. By appropriately specifying the hierarchical model, estimates for these combinations could still be obtained, because they draw from the full model structure. We have identified that different growth forms have different cover distributions. Our empirical evidence strongly suggests that in plots where there are many small entities from the same growth form, such as for forbs and grasses, the cumulative cover of that growth form (when derived from transformations of ordinal data) may amplify and inaccurately describe the structural complexity of vegetation communities. Identifying and accounting for these distributions in other grouped entities has the potential to further improve summed cover estimates.

We also note potential observer bias for cover estimates of 5%. We acknowledge that visual estimates of cover and counts are subject to inter- and intra-operator error and bias (see Morrison 2016 for review) and this may account for the data digressions from a smooth shaped abundance curve. This is no doubt an artefact of observer preference for regular intervals when estimating cover, rather than a true representation of plant cover. In our case study analyses, the high frequency of estimates of 5% cover required a split-model approach where cover data were treated in two separate models. Given this right-skewed distribution and potential bias among disparate datasets (Appendix S3—Figures 1-4), we propose the split-model approach may serve wider applications. Simulated beta distribution data may not be entirely appropriate when using visually-estimated cover data, but may be useful where other less subjective methods for estimating cover are used (such as point-intercept methods). Given that visual estimates of cover-abundance are the assessment protocol for many floristic surveys, our approach offers a way these data can still be transformed and used with greater confidence, despite the underlying variability and bias. The approach we outline here can rapidly generate robust and defensible transformation estimates that are less prone to inflating summed cover estimates.

We envisage that our method may be useful when combined with emerging technologies such as 3-dimensional LiDAR or radar sensors that can penetrate vegetation canopies and assess complex structural elements. Furthermore, where large-scale biodiversity assessments, that rely on terrestrial vegetation as indicators of change, seek to integrate site observations to validate or train imagery, vegetation cover data collected in an ordinal scale will be of little benefit. Summation of midpoint transformations have been widely used in remote-sensing applications, no doubt often resulting in overinflated cover estimates. While transformations derived from a beta distribution will reduce the problem of over inflation, summed cover estimates can still exceed 100%. Jennings *et al.* (2009) offer an approach to rescaling summed cover so that cover estimates do not exceed 100% and their approach may be useful where site-based data are integrated to inform remote sensing applications.

Our approach to transforming ordinal estimates of cover using a beta distribution can extend the application of these data beyond the realm of vegetation classification and can salvage information from many millions of floristic records. We expect most large repositories of floristic data will contain cover estimates with multifarious and nuanced ordinal scales. Here we provide a method that can be applied to floristic data in different ordinal scales for transforming and integrating datasets with much greater confidence. We have demonstrated a pan-continental approach to transforming ordinal cover estimates needed to build robust and accurate aggregated cover estimates. We foresee this approach supporting the synthesis of multiple datasets containing legacy data collected in different ordinal scales, especially where the aggregate properties of vegetation cover for different plant groups are of interest. These transformations and the resultant aggregated properties of cover data can support a multitude of uses in ecology from site-scaled, to landscape-scaled and for global applications.

## Supporting information

SupMat

## Acknowledgments

We thank Stephan Hennekens and Borja Jiménez-Alfaro for assisting with extracting information from the sPlot v2.1 database; Michael Lee and Elizabeth Shrader provided contact information for ecologists in West Virginia and we thank Jim Vanderhorst who provided additional details and advice needed to interpret the West Virginia NHP validation data. Wade Blanchart provided advice on the presentation of OP scatterplots and Terry Koen provided advice on scientific rigour. Thanks to Chris Watson, Jian Yen and Samantha Travers for their helpful comments on draft manuscripts. We thank János Podani and two anonymous referees for valuable and insightful comments.

## Author Contribution Statement

- conceived the ideas and designed methodology – ALL
- extracted the case study and validation data – MM
- designed predictive model – JD
- analysed and interpreted the data – ALL
- led the writing of the manuscript – MM
- contributed critically to drafting and revising the manuscript and gave final approval for publication – ALL

## Abbreviations

BBCA: Braun-Blanquet cover-abundance

## References

Abelleira Martínez, O.J., Fremier, A.K., Günter, S., Ramos Bendaña, Z., Vierling, L., Galbraith, S.M., … Ordoñez, J.C. (2016). Scaling up functional traits for ecosystem services with remote sensing: concepts and methods. Ecol Evol 6: 4359–4371.

Braun-Blanquet, J. (1964). Pflanzensoziologie: Grundzüge der Vegetationskunde. 3rd ed. Springer, Wien, New York.

Braun-Blanquet, J. (1932). Plant Sociology: The Study of Plant Communities: Authorized English Translation of Pflanzensoziologie. McGraw Hill.

Bulmer, M.G. (1979). Principles of statistics. 3rd ed. Constable and Company Ltd., United Kingdom.

Cawsey, E.M., Austin, M.P. & Baker, B.L. (2002). Regional vegetation mapping in Australia: a case study in the practical use of statistical modelling. Biodiversity and Conservation 11: 2239–2274.

Chen, J., Shiyomi, M., Bonham, C.D., Yasuda, T., Hori, Y. & Yamamura, Y. (2008a). Plant cover estimation based on the beta distribution in grassland vegetation. Ecological Research 23: 813–819.

Chen, J., Shiyomi, M., Hori, Y. & Yamamura, Y. (2008b). Frequency distribution models for spatial patterns of vegetation abundance. Ecological Modelling 211: 403–410.

Chiarucci, A., Wilson, J.B., Anderson, B.J. & De Dominicis, V. (1999). Cover versus biomass as an estimate of species abundance: does it make a difference to the conclusions? Journal of Vegetation Science 10: 35–42.

Chytrý, M., Hennekens, S.M., Jiménez-Alfaro, B., Knollová, I., Dengler, J., Jansen, F., … Yamalov, S. (2016). European Vegetation Archive (EVA): an integrated database of European vegetation plots. Applied Vegetation Science 19: 173–180.

Cribari-Neto, F. & Zeileis, A. (2010). Beta regression in R. Journal of Statistical Software 34: 1–24.

Damgaard, C. (2014). Estimating mean plant cover from different types of cover data: a coherent statistical framework. Ecosphere 5: 1–7.

Damgaard, C. (2013). Hierarchical and spatially aggregated plant cover data. Ecological Informatics 18: 35–39.

Damgaard, C. (2009). On the distribution of plant abundance data. Ecological Informatics 4: 76–82.

Dengler, J., Jansen, F., Glöckler, F., Peet, R.K., De Cáceres, M., Chytrý, M., … Spencer, N. (2011). The global index of vegetation-plot databases (GIVD): a new resource for vegetation science. Journal of Vegetation Science 22: 582–597.

Doane, D.P. & Seward, L.E. (2011). Measuring skewness: a forgotten statistic? Journal of Statistics Education 19: 1–18.

Faber-Langendoen, D., Aaseng, N., Hop, K., Lew-Smith, M. & Drake, J. (2007). Vegetation classification, mapping, and monitoring at Voyageurs National Park, Minnesota: an application of the U.S. National Vegetation Classification. Applied Vegetation Science 10: 361–374.

Ferrari, S. & Cribari-Neto, F. (2004). Beta regression for modelling rates and proportions. Journal of Applied Statistics 31: 799–815.

Fill, J.M., Forsyth, G.G., Kritzinger-Klopper, S., Le Maitre, D.C. & van Wilgen, B.W. (2017). An assessment of the effectiveness of a long-term ecosystem restoration project in a fynbos shrubland catchment in South Africa. Journal of Environmental Management 185: 1–10.

Guisan, A. & Harrell, F.E. (2000). Ordinal response regression models in ecology. Journal of Vegetation Science 11: 617–626.

Herpigny, B. & Gosselin, F. (2015). Analyzing plant cover class data quantitatively: customized zero-inflated cumulative beta distributions show promising results. Ecological Informatics 26: 18–26.

Irvine, K.M., Rodhouse, T.J. & Keren, I.N. (2016). Extending ordinal regression with a latent zero-augmented beta distribution. Journal of Agricultural, Biological and Environmental Statistics 21: 619–640.

Jennings, M.D., Faber-Langendoen, D., Loucks, O.L., Peet, R.K. & Roberts, D. (2009). Standards for associations and alliances of the U.S. National Vegetation Classification. Ecological Monographs 79: 173–199.

Joanes, D.N. & Gill, C.A. (1998). Comparing measures of sample skewness and kurtosis. Journal of the Royal Statistical Society: Series D (The Statistician) 47: 183–189.

Keith, D. (2004). Ocean Shores to Desert Dunes: The Native Vegetation of New South Wales and the ACT. Department of Environment and Conservation NSW, Sydney.

Knapp, E.E. & Ritchie, M.W. (2016). Response of understory vegetation to salvage logging following a high-severity wildfire. Ecosphere 7: e01550.

Lyons, M.B., Keith, D.A., Warton, D.I., Somerville, M. & Kingsford, R.T. (2016). Model-based assessment of ecological community classifications. Journal of Vegetation Science 27: 704–715.

MacArthur, R. (1960). On the relative abundance of species. The American Naturalist 94: 25–36.

McElhinny, C., Gibbons, P. & Brack, C. (2006). An objective and quantitative methodology for constructing an index of stand structural complexity. Forest Ecology and Management 235: 54–71.

McGill, B.J., Etienne, R.S., Gray, J.S., Alonso, D., Anderson, M.J., Benecha, H.K., … White, E.P. (2007). Species abundance distributions: moving beyond single prediction theories to integration within an ecological framework. Ecology Letters 10: 995–1015.

Meyer, D., Dimitriadou, E., Hornik, K., Weingessel, A., Leisch, F. & (2017). e1071: misc functions of the Department of Statistics, Probability Theory Group. In. R package version 1.6-8, TU Wien.

Morlon, H., White, E.P., Etienne, R.S., Green, J.L., Ostling, A., Alonso, D., … Zillio, T. (2009). Taking species abundance distributions beyond individuals. Ecology Letters 12: 488–501.

Morrison, L.W. (2016). Observer error in vegetation surveys: a review. Journal of Plant Ecology 9: 367–379.

Mueller-Dombois, D. & Ellenberg, H. (1974). Aims and Methods of Vegetation Ecology. Wiley New York, NY.

Peet, R.K., Lee, M.T., Jennings, M.D. & Faber-Langendoen, D. (2013). VegBank: The vegetation plot archive of the Ecological Society of America. In, http://vegbank.org.

Pereira, H.M., Ferrier, S., Walters, M., Geller, G.N., Jongman, R.H.G., Scholes, R.J., … Wegmann, M. (2013). Essential biodiversity variables. Science 339: 277–278.

Pereira, H.M., Leadley, P.W., Proença, V., Alkemade, R., Scharlemann, J.P.W., Fernandez-Manjarrés, J.F., … Walpole, M. (2010). Scenarios for global biodiversity in the 21st century. Science 330: 1496–1501.

Pettorelli, N., Laurance, W.F., O’Brien, T.G., Wegmann, M., Nagendra, H. & Turner, W. (2014). Satellite remote sensing for applied ecologists: opportunities and challenges. Journal of Applied Ecology 51: 839–848.

Podani, J. (2006). Braun-Blanquet’s legacy and data analysis in vegetation science. Journal of Vegetation Science 17: 113–117.

Podani, J. (2005). Multivariate exploratory analysis of ordinal data in ecology: pitfalls, problems and solutions. Journal of Vegetation Science 16: 497–510.

R Core Team (2018). R: A language and environment for statistical computing. In. R Foundation for Statistical Computing, Vienna, Austria

Schaminée, J.H., Hennekens, S.M., Chytrý, M. & Rodwell, J.S. (2009). Vegetation-plot data and databases in Europe: an overview. Preslia 81: 173–185.

Schaminée, J.H.J., Janssen, J.A.M., Hennekens, S.M. & Ozinga, W.A. (2011). Large vegetation databases and information systems: new instruments for ecological research, nature conservation, and policy making. Plant Biosystems - An International Journal Dealing with all Aspects of Plant Biology 145: 85–90.

Scholes, R.J. & Biggs, R. (2005). A biodiversity intactness index. Nature 434: 45–49.

Scott, J.J. & Kirkpatrick, J.B. (2008). Rabbits, landslips and vegetation change on the coastal slopes of subantarctic Macquarie Island, 1980–2007: implications for management. Polar Biology 31: 409–419.

Su, Y. & Yajima, M. (2015). R2jags: Using R to run ‘JAGS’. R package version 0.5–7. In, CRAN. R-project.org/package=R2jags.

Tüxen, R. & Ellenberg, H. (1937). Der systematische und ökologische Gruppenwert. Ein Beitrag zur Begriffsbildung und Methodik der Pflanzensoziologie. Mitt. Flor.-Soz. Arbeitsgem 3: 171–184.

van der Maarel, E. (2007). Transformation of cover-abundance values for appropriate numerical treatment: alternatives to the proposals by Podani. Journal of Vegetation Science 18: 767–770.

van der Maarel, E. (1979). Transformation of cover-abundance values in phytosociology and its effects on community similarity. Vegetatio 39: 97–114.

Vanderhorst, J., Byers, E. & Streets, B. (2012). Natural Heritage Vegetation Database for West Virginia. In: Jürgen Dengler, Jens Oldeland, Florian Jansen, Milan Chytrý, Jörg Ewald, Manfred Finckh, Falko Glöckler, Gabriela Lopez-Gonzalez, Robert K Peet & Joop HJ Schaminée (eds.) Vegetation Databases for the 21st Century - 9th Meeting on Vegetation Databases, pp. 440–440. Biodiversity, Evolution and Ecology of Plants (BEE) Biocentre Klein Flottbek and Botanical Garden, University of Hamburg, Hamburg.

Warton, D.I. & Hui, F.K.C. (2011). The arcsine is asinine: the analysis of proportions in ecology. Ecology 92: 3–10.

Whittaker, R.H. (1965). Dominance and diversity in land plant communities. Science 147: 250–260.

Wiser, S.K. & De Cáceres, M. (2013). Updating vegetation classifications: an example with New Zealand’s woody vegetation. Journal of Vegetation Science 24: 80–93.

Zuur, A.F., Hilbe, J.M. & Ieno, E.N. (2013). A beginner’s guide to GLM and GLMM with R: a frequentist and Bayesian perspective for ecologists. Highland Statistics Ltd, Newburgh, United Kingdom.

