## Supplementary material for "Species abundance distributions should underpin ordinal cover-abundance transformations": SupMat

**Appendices**

This supplementary material supports the paper McNellie, M.J., Dorrough, J and Oliver, I. Skewed species abundance should underpin ordinal cover-abundance transformation.

**Authors**:

<https://orcid.org/0000-0002-8162-2551>

Josh Dorrough^3^

<https://orcid.org/0000-0001-7099-1442>

Ian Oliver^4, 5^

<https://orcid.org/0000-0003-3276-6644>

^1^ Fenner School of Environment and Society, The Australian National University, Canberra, ACT, Australia

^2^ Office of Environment and Heritage New South Wales, Wagga Wagga, NSW, Australia

^3^ Office of Environment and Heritage New South Wales, Merimbula, NSW, Australia

^4^ School of Environmental and Rural Sciences, University of New England, Armidale, NSW, Australia

^5^ Office of Environment and Heritage, Gosford, NSW, Australia

**Table of Contents**

Appendix S1: Sample R code to build Bayesian hierarchical mixed-effects model to estimate predicted means (PM) for each growth form for BBCA3 – BBCA

Appendix S2 Figure 1: Geographic spread of 2 809 plots used to compare visual quantitative cover estimates of 95 812 floristic records with a transformed Braun–Blanquet cover–abundance (BBCA) scale.

Appendix S3 Figure 1: Floristics data sourced from Waterton Lakes National Park, British Columbia, Canada.

Appendix S3 Figure 2: Floristics data sourced from Hawai‘i Volcanoes National Park, Hawai‘i Island.

Appendix S3 Figure 3: Floristics data sourced from Fire Island National Park, Long Island, New York.

Appendix S3 Figure 4: Floristics data sourced from Great Britain Countryside Survey (1990), England, Scotland and Wales.

Appendix S4 Table 1: Results from linear regressions with zero-intercepts using observed and predicted summed cover for six plant growth forms and total summed cover transformed using either the predicted mean or historical transformations. For each linear regression the fit of the observed and predicted summed cover are assessed using root mean squared deviation (RMSD), slope (with 95% confidence intervals) and adjusted R^2^.

Appendix S5 Figure 1: Scatter plots of case study dataset showing the relationships between summed cover for the visual estimates of cover (0.1%–100%) compared to the sum of cover when transformed using predicted mean (PM) for different growth forms

Appendix S5 Figure 2: Scatter plots of case study dataset showing the relationships between summed cover for the visual estimates of cover (0.1%–100%) compared to the sum of cover when transformed using Tüxen and Ellenberg (1937) (T & E) for different growth forms

Appendix S5 Figure 3: Scatter plots of case study dataset showing the relationships between summed cover for the visual estimates of cover (0.1%–100%) compared to the sum of cover when transformed using Braun-Blanquet (1964) (BB) for different growth forms

Appendix S5 Figure 4: Scatter plots of case study dataset showing the relationships between summed cover for the visual estimates of cover (0.1%–100%) compared to the sum of cover when transformed using van der Maarel (2007) (vdM) for different growth forms

Appendix S6 Preparation of the validation dataset and results of linear regression and RMSD analyses.

Appendix S6 - Table 1: Results from linear regressions with zero-intercepts using validation dataset of observed and predicted summed cover for six plant growth forms and total summed cover transformed using either the predicted mean or Tüxen & Ellenberg (1937) transformations. For each linear regression the fit of the observed and predicted summed cover are assessed using root mean squared deviation (RMSD), slope (with 95% confidence intervals) and adjusted R^2^.

Appendix S7 Figure 1: Scatter plots of validation data showing the relationships between summed cover for the visual estimates of cover (0.01%–100%) compared to the sum of cover when transformed by predicted mean (PM) for different growth forms

Appendix S7 Figure 2: Scatter plots of validation data showing the relationships between summed cover for the visual estimates of cover (0.01%–100%) compared to the sum of cover when transformed by Tüxen and Ellenberg (1937) (T & E) for different growth forms

Appendix S1: Sample R code to build Bayesian hierarchical mixed-effects model to estimate predicted means (PM) for each growth form for BBCA3 – BBCA6. This needs to be slightly modified to estimate values for classes BBCA1 & BBCA2 or to obtain values without growth form. Dataset can be found at <https://figshare.com/s/0681b3b14d8c8402b068>.

```
data<- read.csv(file = "bbca.csv") #data frame of visual cover , site , species , BBCA bands 1-6 (band)  and GF (growth form)
#bands 3-6
	bbca3_6<-subset(data, band>=3)
	bbca3_6$fBand <- factor(bbca3_6$band) #BBCA bands 3,4,5,6 as factor

#transform visual % and scale between 0-1
n<-nrow(bbca3_6)-1
	p<-( bbca3_6$Visual-5)/(100-5)
		bbca3_6$PropCover <- (p*n +0.5)/(n+1)
#Interaction model
X <- model.matrix(~ 1+GF * fBand, data = bbca3_6) #model matrix for the interaction between GF and Band
K <- ncol(X)  		#Number of columns
head(X)
nrow(X)

#Random effects:
Site <- as.numeric(as.factor(bbca3_6$fSite))
Site
Nre <- length(unique(bbca3_6$fSite))
Nre
win.data <- list(Y     = bbca3_6$PropCover, 	#response
                 X     = X,               	#fixed effects
                 N     = nrow(bbca3_6),
                 K     = ncol(X),
                 Site  = Site, 		#site random effect
                 Nre  = Nre) #number of levels of the random effect

win.data

sink("bbca3_6.txt")		#JAGs code, stored as separate text file, identical pasted below
cat("
    model{
    #1A. Priors beta and sigma
    for (i in 1:K) { beta[i] ~ dnorm(0, 0.001)}
    theta ~ dunif(0, 100) #sd of 1000 used for bands 1 &2
    #1B. Priors random plot effects and sigma_Site
    for (i in 1:Nre) { a[i] ~ dnorm(0, tau1_Site)}
    sigma1_Site ~ dunif(0, 10)
    tau1_Site <- 1 / (sigma1_Site * sigma1_Site)
    #2. Likelihood
    for (i in 1:N) {
    Y[i]          ~ dbeta(shape1[i], shape2[i])# shape parameters for the beta distribution
    shape1[i] <- theta * pi[i]
    shape2[i] <- theta * (1 - pi[i])
    logit(pi[i]) <- mu[i]
    eta[i]       <- inprod(beta[], X[i,])
    mu[i]        <- eta[i] + a[Site[i]]
    }
    }
    ",fill = TRUE)
sink()
#initial values
inits  <- function () {
  list(
    beta       = rnorm(ncol(X), 0, 0.001),
    theta = runif(0, 100), #sd of 1000 used for bands 1 & 2
    a          = rnorm(Nre, 0, 0.1),
    sigma1_Site = runif(1, 0, 10)
  )  }

params <- c("beta", "a" ,"sigma1_Site","theta") # save beta's, a,sigma site and theta

#preliminary run
beta3 <- jags(data       = win.data,
              inits      = inits,
              parameters = params,
              model      = "transtest.txt",
              n.thin     = 15,
              n.chains   = 3,
              n.burnin   = 3000,
              n.iter     = 5000)

#update
beta3.1  <- update(beta3, n.iter = 15000, n.thin = 15)
beta3.out <- beta3.1$BUGSoutput		# save output

#print (beta3.out, digits = 3)
K <- ncol(X)
beta3_OUTV3 <- MyBUGSOutput(beta3.out,
                            c(uNames("beta", K), "sigma1_Site", "theta"),
                            VarNames  = MyNames)
print(beta3_OUTV3, digits =5)

#extract betas for each iteration in each chain
Beta3.mcmc <- beta3.out$sims.list$beta

#obtain predicted means and credible intervals
NewTrans <- expand.grid(GF= levels(bbca3_6$GF),
                        fBand        = levels(bbca3_6$fBand))
Xp     <- model.matrix(~  GF*fBand,
                       data = NewTrans)
mu <- Xp %*% t(Beta3.mcmc)
mu1 <- (exp(mu) / (1 + exp(mu)))
L <- GetCIs(mu1)
L
predicted_means_BBCA3_6 <- cbind(NewTrans,L)
```

**Appendix S2**

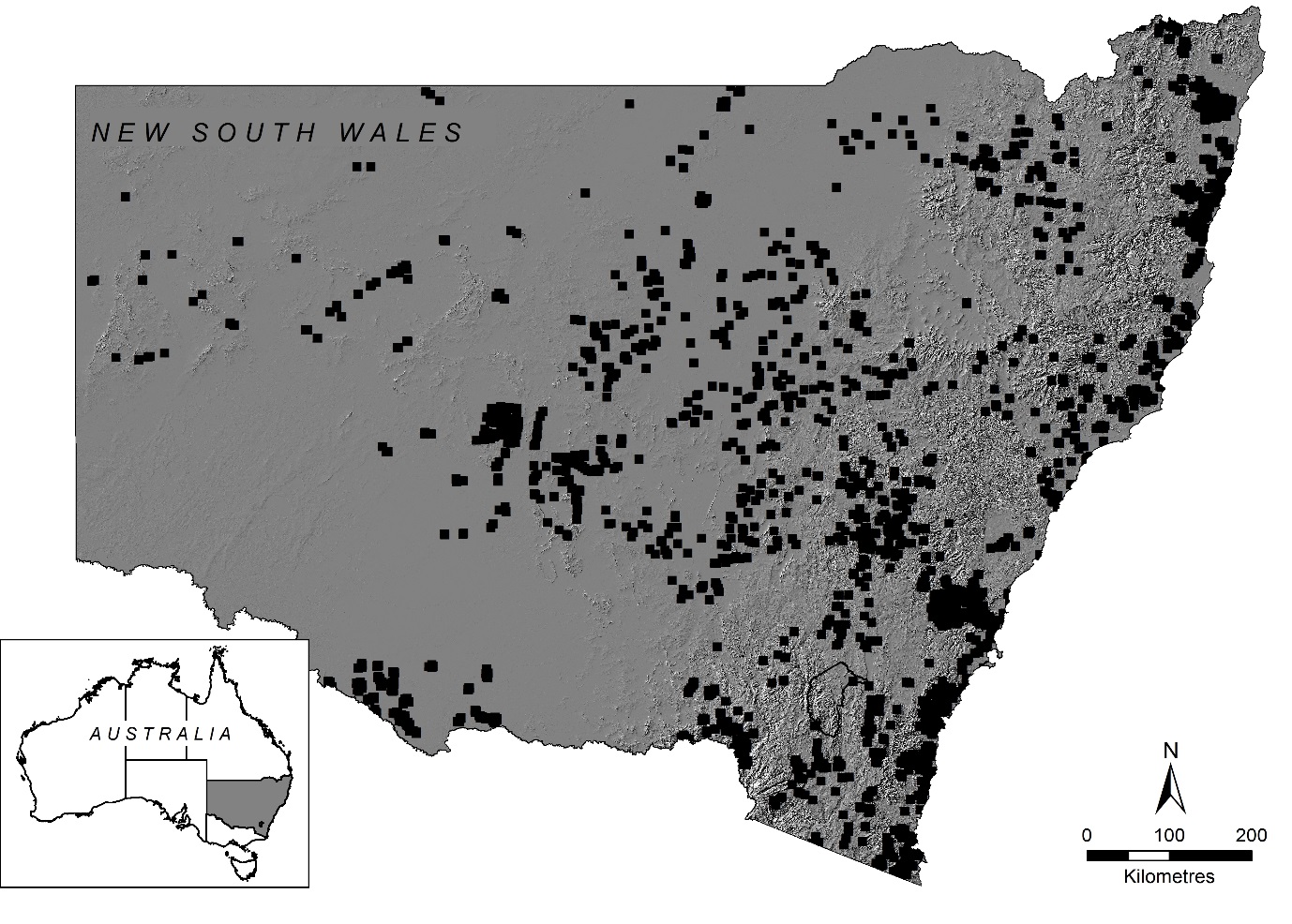

### Appendix S2 Figure 1: Geographic spread of 2 809 plots used to compare visual quantitative cover estimates of 95 812 floristic records with a transformed Braun–Blanquet cover–abundance (BBCA) scale.

**Appendix S3**

The right-skewed distribution of plant cover is not unique to our dataset. We sourced data from unrestricted, free, digital repositories where cover estimates were recorded as quantitative estimates. We were able to find suitable datasets from Vegetation Mapping Inventory Projects housed by the National Park Service (U.S. Department of the Interior) <https://science.nature.nps.gov/im/inventory/veg/products.cfm> and selected datasets from Canada and two US States (Hawai‘i and New York). We also used the Countryside Survey of Great Britain (https://catalogue.ceh.ac.uk/eidc/documents). We applied the same decision rules and allocated visual field-based cover estimates to their equivalent 1–6 Braun-Blanquet cover-abundance (BBCA) class (see Table 1 in manuscript). We found these datasets showed similarly positively skewed distributions with 70–80% of observations having cover less than 5%.

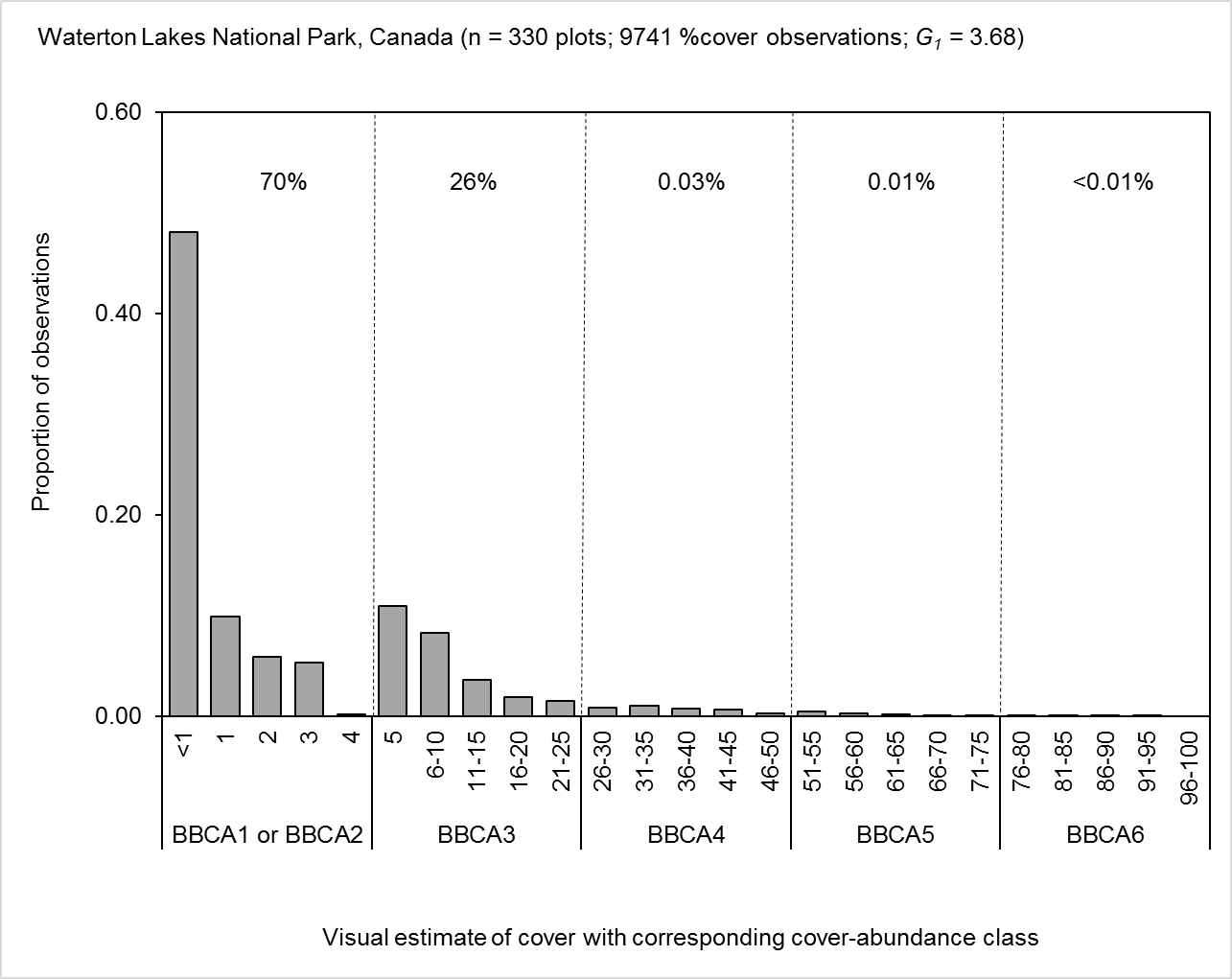
Appendix S3 Figure 1: Floristics data sourced from Waterton Lakes National Park, British Columbia, Canada.

Data sourced from Field Data Microsoft Access Database at <https://science.nature.nps.gov/im/inventory/veg/project.cfm?ReferenceCode=1047714> on 12^th^ July 2017. These data were surveyed for the Vegetation Mapping Inventory Project for Glacier National Park (Hop et al. 2007) and analyses are based on only Waterton Lakes National Park plots (n = 330 plots containing 9 741 observations of cover). Continuous cover estimates ranged from 0.5–95. Abundance records were not included in the data supplied, therefore BBCA1 and BBCA2 (both less than 5% cover) are combined. Plot area = 400 m^2^ (Hop et al. 2007). Adjusted Fisher-Pearson skewness coefficient (*G_1_*) = 3.68. Dashed vertical lines show cut points between each BBCA class. Numbers between the dashed lines show the proportion of each class in the dataset.

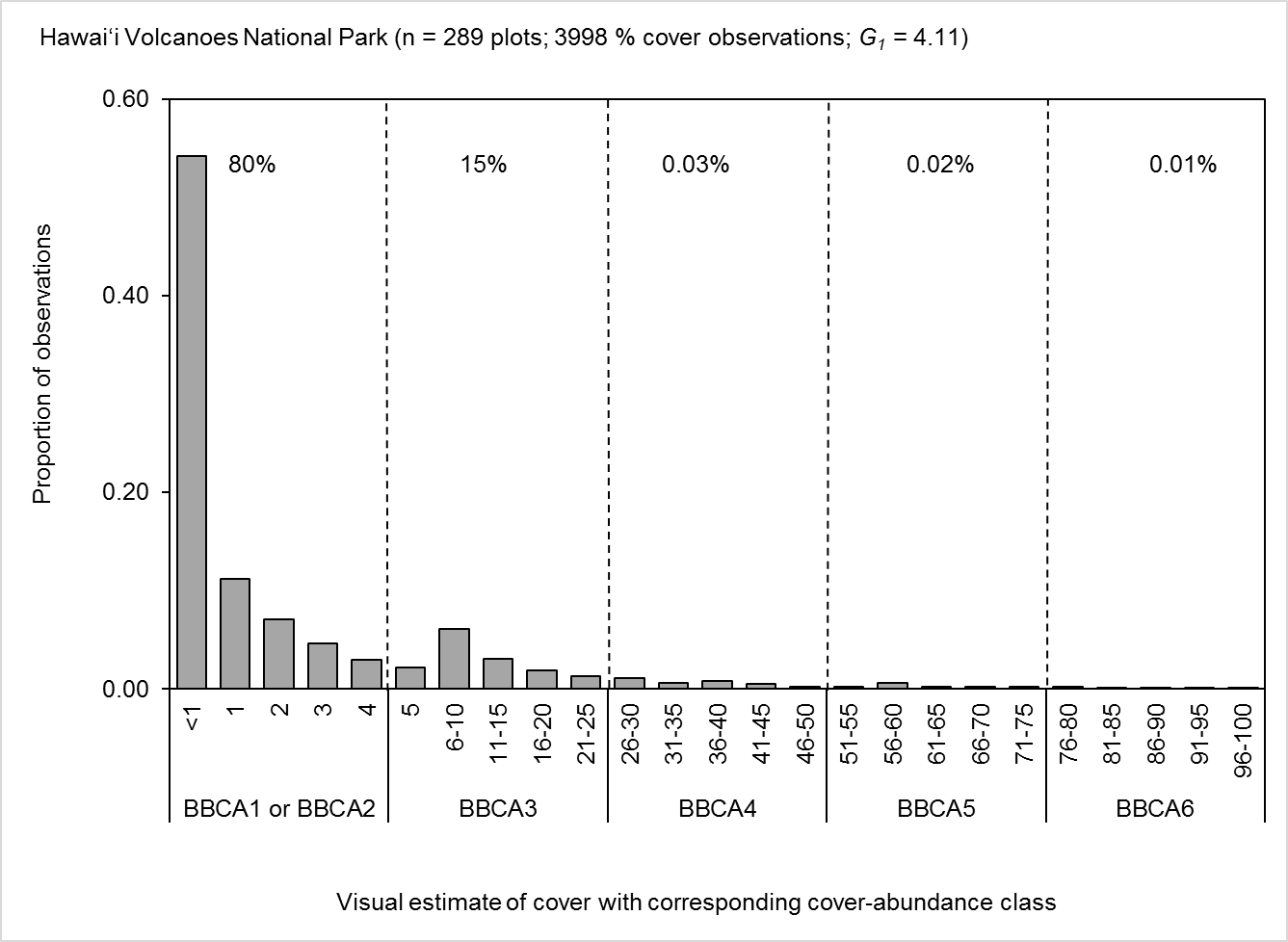
Appendix S3 Figure 2: Floristics data sourced from Hawai‘i Volcanoes National Park, Hawai‘i Island.

Data sourced from <https://science.nature.nps.gov/im/inventory/veg/project.cfm?ReferenceCode=2230298> on 12^th^ July 2017. These data were surveyed for the Vegetation Mapping Inventory Project for Hawai‘i Volcanoes National Park (Green et al. 2015) and analyses are based on only ‘classification’ plots (n = 289 plots containing 3 998 observations of cover). Continuous cover estimates ranged from 0.5–98. Abundance records were not included in the data supplied, therefore BBCA1 and BBCA2 (both less than 5% cover) are combined. Plot area = 400 m^2^ (Green et al. 2015). Adjusted Fisher-Pearson skewness coefficient (*G_1_*) = 4.11. Dashed vertical lines show cut points between each BBCA class. Numbers between the dashed lines show the proportion of each class in the dataset.

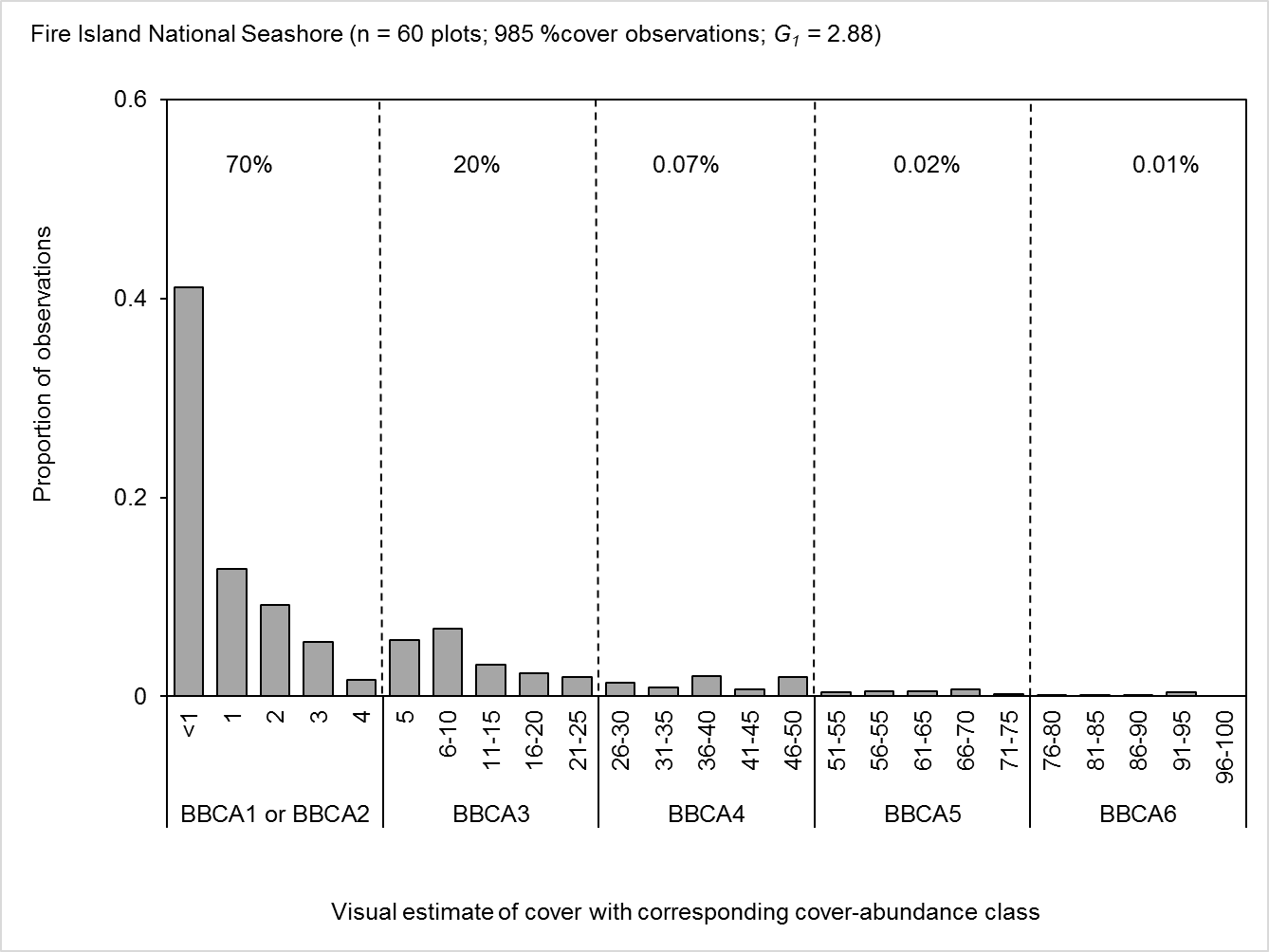
Appendix S3 Figure 3: Floristics data sourced from Fire Island National Park, Long Island, New York.

Data sourced from <https://irma.nps.gov/DataStore/Reference/Profile/2233594> on 12th July 2017. These data were surveyed for the Vegetation Mapping Inventory Project for Fire Island National Seashore (Klopfer et al. 2002) and analyses are based on only ‘classification’ plots (n = 66 plots containing 985 observations of cover). Continuous cover estimates ranged from 0.5–95. Abundance records were not included in the data supplied, therefore BBCA1 and BBCA2 (both less than 5% cover) are combined. Plot area = 400 m^2^ (Klopfer et al. 2002). Adjusted Fisher-Pearson skewness coefficient (*G_1_*) = 2.88. Dashed vertical lines show cut points between each BBCA class. Numbers between the dashed lines show the proportion of each class in the dataset.

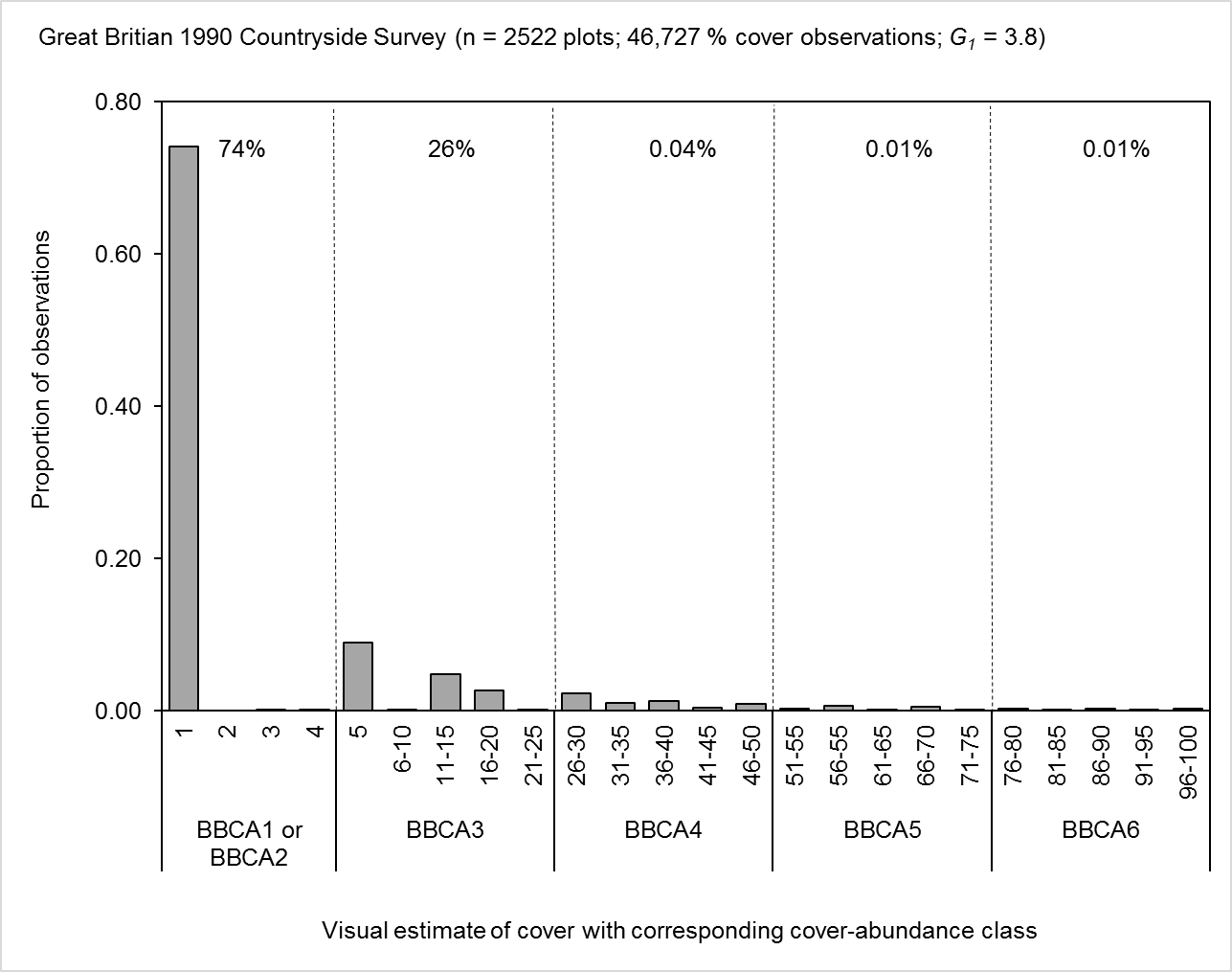
Appendix S3 Figure 4: Floristics data sourced from Great Britain Countryside Survey (1990), England, Scotland and Wales.

Data sourced from <https://doi.org/10.5285/26e79792-5ffc-4116-9ac7-72193dd7f191> on 11^th^ July 2017. This resource is available under [the Open Government Licence (OGL)](http://www.nationalarchives.gov.uk/doc/open-government-licence). Contains data supplied by Natural Environment Research Council. © NERC (Centre for Ecology & Hydrology). These data were surveyed for the 1990 Countryside Survey (of England, Scotland and Wales) and analyses are based on plots coded with ‘X’ in the dataset (n = 2 522 plots containing 46 727 observations of cover). Cover was recorded in 1% increments up to 5% and in 5% increments thereafter. Observations ranged from 1–100%. Abundance records were not included in the data supplied, therefore BBCA1 and BBCA2 (both less than 5% cover) are combined. Plot area = 200 m^2^ (Barr et al. 1990). Adjusted Fisher-Pearson skewness coefficient (*G_1_*) = 3.8.

**Supplementary Material References**

Barr, C.J., Bunce, R.G.H.; Gillespie, M.K.; Hallam, C.J.; Howard, D.C.; Maskell, L.C.; Ness, M.J.; Norton, L.R.; Scott, R.J.; Smart, S.M.; Stuart, R.C.; Wood, C.M. 2014. Countryside Survey 1990 vegetation plot data. NERC Environmental Information Data Centre.<https://doi.org/10.5285/26e79792-5ffc-4116-9ac7-72193dd7f191>

Green, K., M. Hall, C. Lopez, A. Ainsworth, M. Selvig, K. Akamine, S. Fugate, K. Schulz, D. Benitez, M. Wasser, and G. Kudray. 2015. Vegetation mapping inventory project: Hawaiʻi Volcanoes National Park. Natural Resource Report NPS/PACN/NRR—2015/966. National Park Service, Fort Collins, Colorado.

Hop, K., M. Reid, J. Dieck, S. Lubinski, and S. Cooper. 2007. U.S. Geological Survey-National Park Service Vegetation Mapping Program: Waterton-Glacier International Peace Park. U.S. Geological Survey, Upper Midwest Environmental Sciences Center, La Crosse, Wisconsin, August 2007. 131 pp. + Appendixes A-L.

Klopfer S.D., Olivero, A., Sneddon, L. 2002. Final Report of the NPS Vegetation Mapping Project at Fire Island National Seashore. Virginia Polytechnic Institute and State University. Blacksburg, VA.

Appendix S4 Table 1: Results from linear regressions with zero-intercepts using case study dataset of observed and predicted summed cover for six plant growth forms and total summed cover transformed using either the predicted mean or historical transformations. For each linear regression the fit of the observed and predicted summed cover are assessed using root mean squared deviation (RMSD), slope (with 95% confidence intervals) and adjusted R^2^used in Figure 3. Number of observations (n) for trees (n = 11 953); shrubs (n = 18 545); grasses (n = 21 575); forbs (n = 30 017); ferns (n = 3 671); other (n = 10 051) and summed total cover (n = 2 809).

| **Growth form** | **Transformation** | **RMSD** | **Slope** | **Lower slope** | **Upper slope** | **Adjusted R^2^** |
| --- | --- | --- | --- | --- | --- | --- |
| Trees | Tüxen & Ellenberg | 13.24 | 0.84 | 0.83 | 0.84 | 0.95 |
|  | Braun-Blanquet | 18.56 | 0.74 | 0.74 | 0.75 | 0.93 |
|  | van der Maarel | 21.4 | 0.7 | 0.69 | 0.71 | 0.93 |
|  | Predicted Mean | 9.5 | 1.03 | 1.02 | 1.04 | 0.95 |
| Shrubs | Tüxen & Ellenberg | 15.01 | 0.67 | 0.66 | 0.68 | 0.87 |
|  | Braun-Blanquet | 9.65 | 0.81 | 0.8 | 0.82 | 0.91 |
|  | van der Maarel | 16.21 | 0.64 | 0.63 | 0.64 | 0.9 |
|  | Predicted Mean | 6.44 | 1.01 | 1 | 1.02 | 0.94 |
| Grasses | Tüxen & Ellenberg | 16.55 | 0.78 | 0.77 | 0.78 | 0.94 |
|  | Braun-Blanquet | 29.41 | 0.62 | 0.61 | 0.63 | 0.9 |
|  | van der Maarel | 23.94 | 0.67 | 0.66 | 0.68 | 0.93 |
|  | Predicted Mean | 9.01 | 1.02 | 1.01 | 1.03 | 0.96 |
| Forbs | Tüxen & Ellenberg | 16.17 | 0.46 | 0.45 | 0.47 | 0.73 |
|  | Braun-Blanquet | 32.7 | 0.26 | 0.25 | 0.27 | 0.61 |
|  | van der Maarel | 20.15 | 0.39 | 0.39 | 0.4 | 0.73 |
|  | Predicted Mean | 5.69 | 0.91 | 0.9 | 0.93 | 0.85 |
| Ferns | Tüxen & Ellenberg | 5.62 | 0.8 | 0.79 | 0.82 | 0.9 |
|  | Braun-Blanquet | 9.02 | 0.66 | 0.64 | 0.67 | 0.84 |
|  | van der Maarel | 7.76 | 0.7 | 0.68 | 0.71 | 0.88 |
|  | Predicted Mean | 3.94 | 1.02 | 1.01 | 1.04 | 0.93 |
| Others | Tüxen & Ellenberg | 6.8 | 0.81 | 0.8 | 0.82 | 0.9 |
|  | Braun-Blanquet | 10.76 | 0.66 | 0.65 | 0.67 | 0.85 |
|  | van der Maarel | 11.27 | 0.63 | 0.62 | 0.64 | 0.89 |
|  | Predicted Mean | 4.79 | 1.05 | 1.03 | 1.06 | 0.93 |
| Total | Tüxen & Ellenberg | 41.47 | 0.74 | 0.74 | 0.75 | 0.96 |
|  | Braun-Blanquet | 79.37 | 0.57 | 0.57 | 0.58 | 0.92 |
|  | van der Maarel | 67.29 | 0.61 | 0.61 | 0.62 | 0.96 |
|  | Predicted Mean | 18.21 | 1.01 | 1.01 | 1.02 | 0.97 |

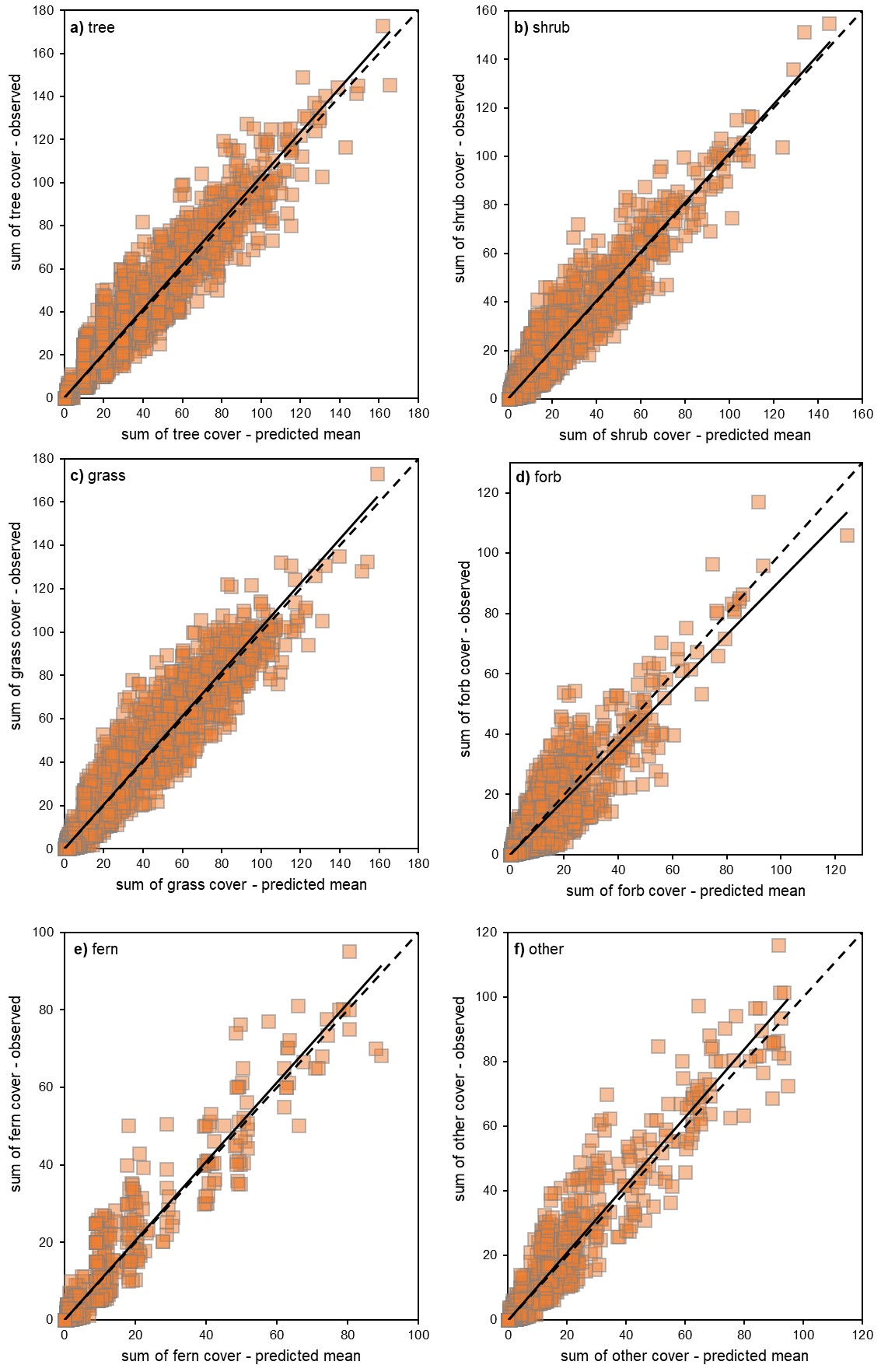
Appendix S5 Figure 1: Scatter plots of case study dataset showing the relationships between summed cover for the visual estimates of cover (0.1%–100%) compared to the sum of cover when transformed using predicted mean (PM) for different growth forms. Number of observations (n) for a) trees (n = 11 953); b) shrubs (n = 18 545); c) grasses (n = 21 575); d) forbs (n = 30 017); e) ferns (n = 3 671) and f) other (n = 10 051). The ideal 1:1 line of best fit is shown as dashed line and solid black line shows slope of the regression through the origin. Transformations that overestimate will have a slope <1.

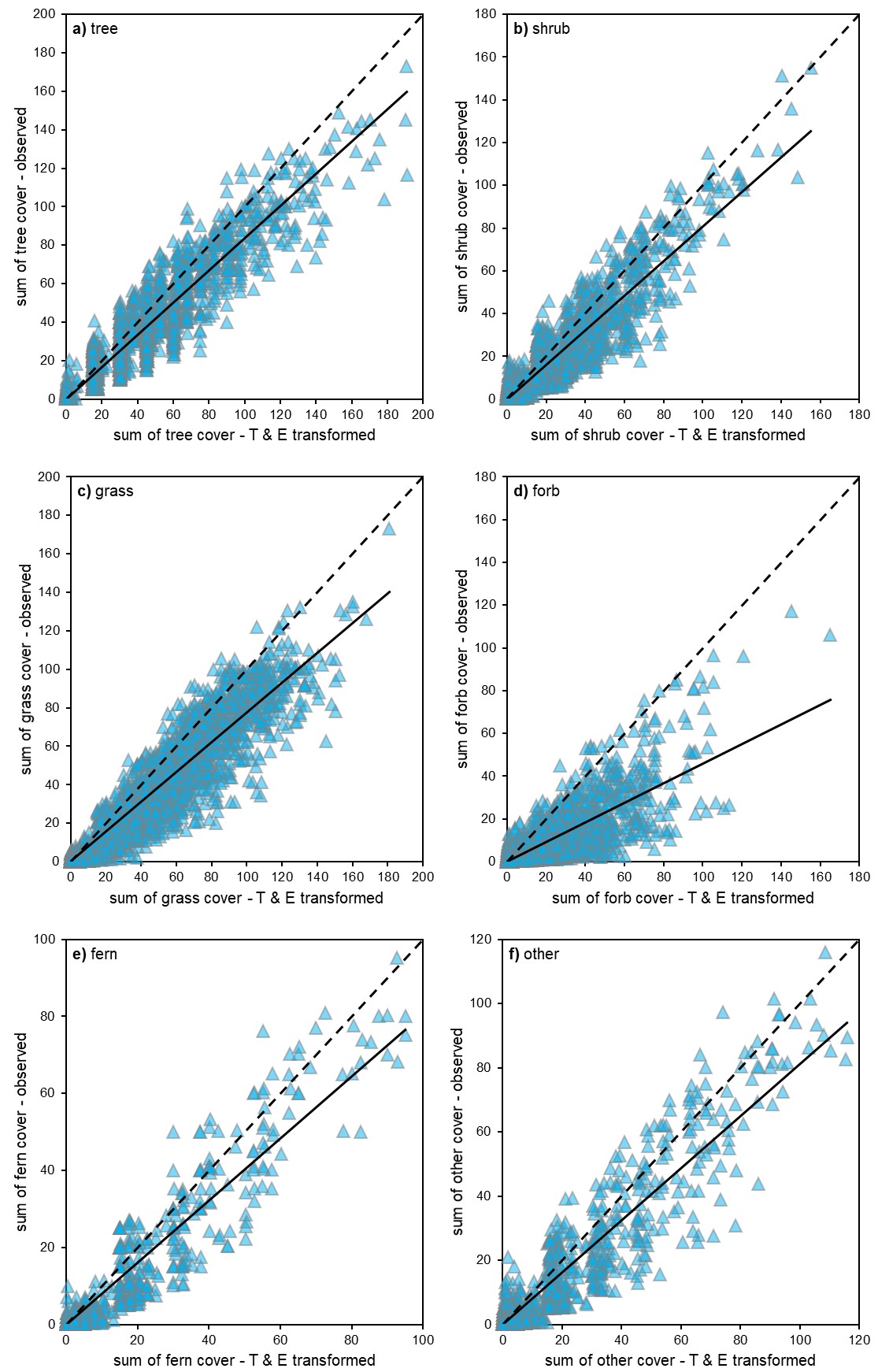
Appendix S5 Figure 2: Scatter plots of case study dataset showing the relationships between summed cover for the visual estimates of cover (0.1%–100%) compared to the sum of cover when transformed by Tüxen and Ellenberg (1937) (T & E) for different growth forms. Number of observations (n) for **a)** trees (n = 11 953); **b)** shrubs (n = 18 545); **c)** grasses (n = 21 575); **d)** forbs (n = 30 017); **e)** ferns (n = 3 671) and **f)** other (n = 10 051). The ideal 1:1 line of best fit is shown as dashed line and solid black line shows slope of the regression through the origin. Transformations that overestimate will have a slope <1.

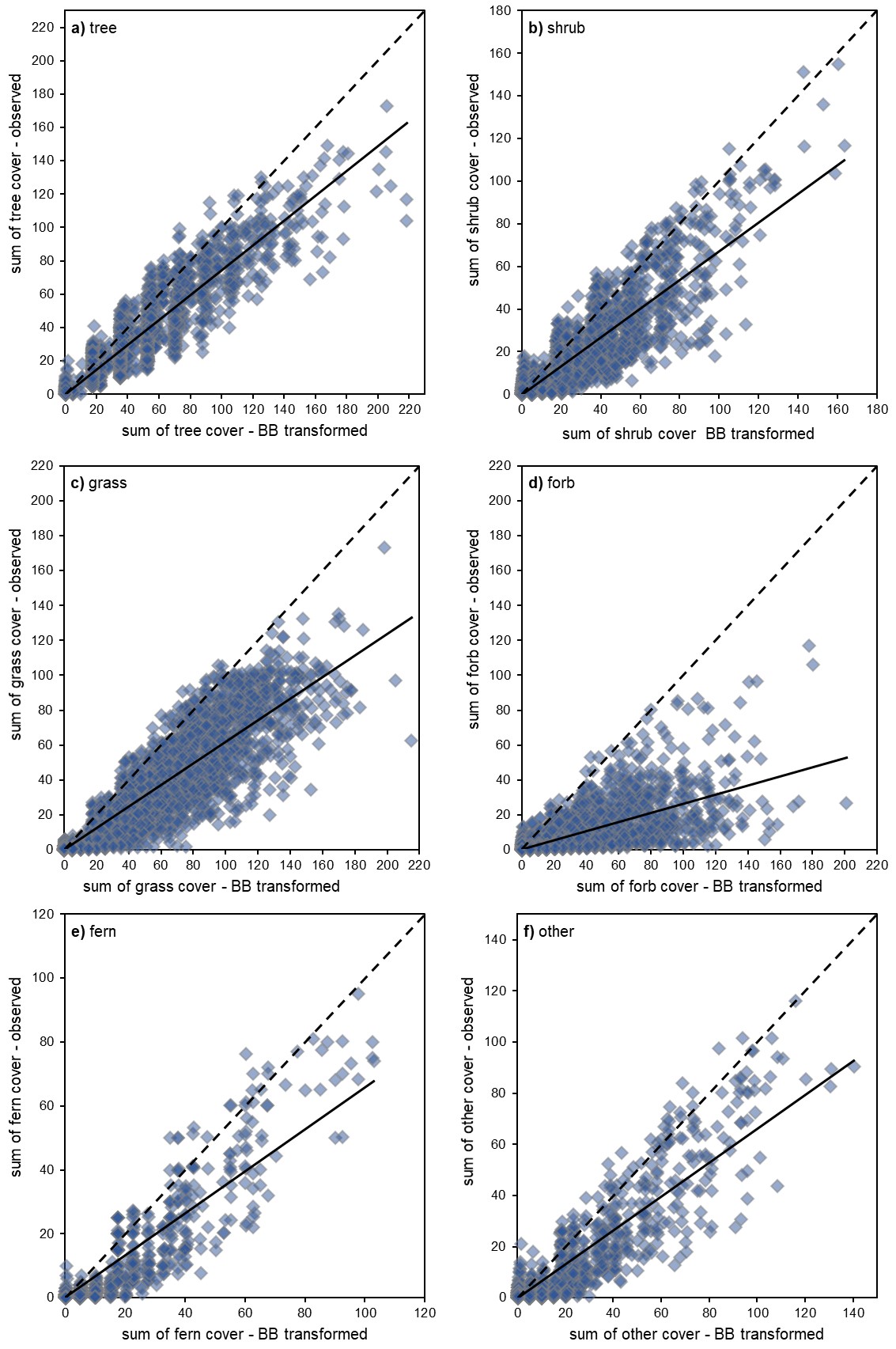
Appendix S5 Figure 3: Scatter plots of case study dataset showing the relationships between summed cover for the visual estimates of cover (0.1%–100%) compared to the sum of cover when transformed by Braun-Blanquet (1964) (BB) for different growth forms. Number of observations (n) for **a**) trees (n = 11 953); **b**) shrubs (n = 18 545); **c**) grasses (n = 21 575); **d**) forbs (n = 30 017); **e**) ferns (n = 3 671) and **f**) other (n = 10 051). The ideal 1:1 line of best fit is shown as dashed line and solid black line shows slope of the regression through the origin. Transformations that overestimate will have a slope <1.

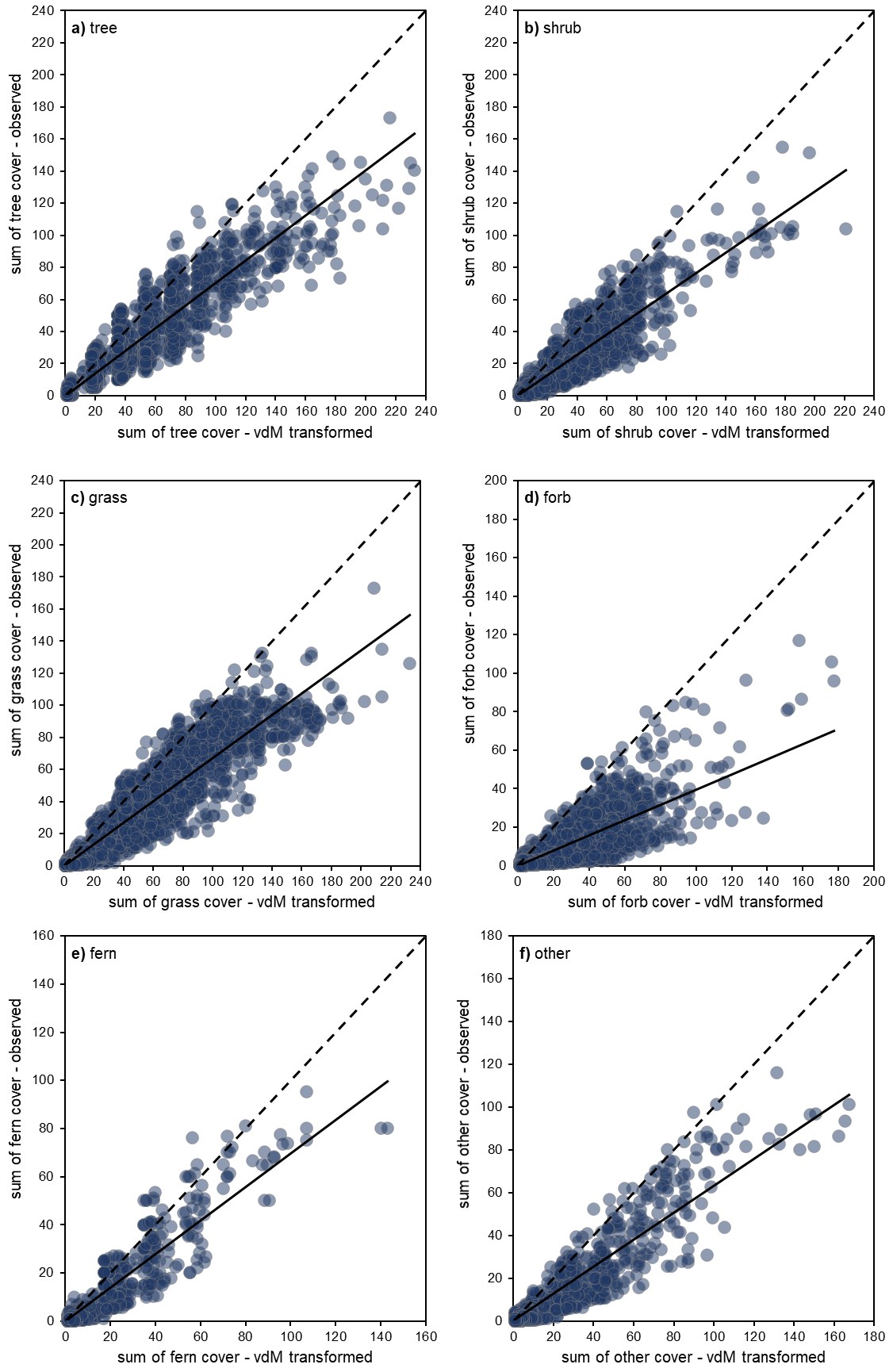
Appendix S5 Figure 4: Scatter plots of case study dataset showing the relationships between summed cover for the visual estimates of cover (0.1%–100%) compared to the sum of cover when transformed by van der Maarel (2007) (vdM) for different growth forms. Number of observations (n) for **a**) trees (n = 11 953); **b**) shrubs (n = 18 545); **c**) grasses (n = 21 575); **d**) forbs (n = 30 017); **e**) ferns (n = 3 671) and **f**) other (n = 10 051). The ideal 1:1 line of best fit is shown as dashed line and solid black line shows slope of the regression through the origin. Transformations that overestimate will have a slope <1.

**Supplementary Material References**

Braun-Blanquet, J. (1964) Pflanzensoziologie: Grundzüge Der Vegetationskunde., 3rd edn. Springer, Wien, New York.

Tüxen, R. & Ellenberg, H. (1937) Der Systematische Und Ökologische Gruppenwert. Ein Beitrag Zur Begriffsbildung Und Methodik Der Pflanzensoziologie. Mitt. Flor.-Soz. . Arbeitsgem, 3, 171-184.

van der Maarel, E. (2007) Transformation of Cover-Abundance Values for Appropriate Numerical Treatment: Alternatives to the Proposals by Podani. Journal of Vegetation Science, 18, 767-770.

### **Appendix S6 Preparation of the validation dataset and results of linear regression and RMSD analyses.**

Whilst VegBank has a primary role for enabling the vegetation classification, large volumes of individual floristic observations are available for synthesis and analysis. VegBank database structure is elegantly simple and accessible for external users. Data are easy to find and download, standardised taxonomic nomenclature and have well-documented metadata. The West Virginia Division of Natural Resources, Natural Heritage Program survey vegetation plots to classify and characterise natural and semi-natural vegetation in the state of West Virginia, primarily to inform conservation (Vanderhorst et al. 2012). A subset of these plots were extracted from VegBank (Peet et al. 2013; accessed 28th Aug 2018) and used as the validation dataset*.*

The validation dataset contained 2 227 plots with 51 497 observations of species level taxa with visual observations of cover for 2 190 species. The subset of data we extracted matched our case study data specifications in that it contained a (i) full floristic inventory surveyed from 400 m^2^ fixed area plots and (ii) visual estimates of plant cover recorded as continuous percentage cover estimates. Where multiple estimates of cover were recorded for the same species within a plot (e.g. assessing species cover within strata), only the maximum cover estimate was used to sum cover (James Vanderhorst pers comm 4^th^ Sep 2018). The minimum observed cover estimate was 0.01% (compared to 0.1% in case study dataset). We note that this much lower estimate of cover may hinder the model performance for smaller growth forms such as forbs. In addition, our validation dataset did not contain counts of individuals which was needed to split observations of less than 5% cover into BBCA1 or BBCA2. To overcome these missing abundance estimates, we replicated the proportions of growth forms in BBCA1 and BBCA2 observed in our case study dataset; ergo 95% of trees; 83% of shrubs; 51% of grasses; 58% of forbs and ferns and 80% of others were randomly allocated to BBCA1. The remaining observations less than 5% cover were allocated to BBCA2.

All taxa were allocated to growth form by matching species name to those listed by Engemann et al. (2016). We split the taxa within the herb group by family so we could validate grass and fern growth forms. Each species was assigned to one of six growth forms using existing allocations (Engemann et al. 2016). The herb growth form was partitioned by the families Poaceae, Cyperaceae, Juncaceae, Liliaceae and Xyridaceae into the grass and grass-like growth forms. Families Dennstaedtiaceae, Dryopteridaceae, Lycopodiaceae, Ophioglossaceae, Osmundaceae, Polypodiaceae and Thelypteridaceae were partitioned into ferns. Vines were classified as other, which would have also included epiphytes and lianas, but none were recorded in the validation species list. Number of observations (n) for trees (n = 13 850); shrubs (n = 4 796); grasses (n = 8 546); forbs (n = 20 911); ferns (n = 2 561) and other (n = 833).

We tested the fit of predicted transformation values for summed growth form cover and total cover to assess the robustness of the PM transformations and evaluate the PM and best performing historical transformation -- Tüxen and Ellenberg (1937) transformations.

**Supplementary Material References**

Vanderhorst, J, Byers, E, Streets, B., 2012. Natural Heritage Vegetation Database for West Virginia. In Dengler, J., Oldeland, J., Jansen, F., Chytry, M., Ewald, J., Fickh, M., . . . Schaminée, J. (2012). Vegetation databases for the 21st century. *Biodiversity and Ecology* 4: 447.

Engemann, K., Sandel, B., Boyle, B., Enquist, B.J., Jørgensen, P.M., Kattge, J., . . . Svenning, J.C., 2016. A plant growth form dataset for the New World. Ecology 97, 3243-3243.

Peet, R.K., M.T. Lee, M.D. Jennings, D. Faber-Langendoen (eds). 2013. VegBank: The vegetation plot archive of the Ecological Society of America. http://vegbank.org; searched on [28 Aug 2018].

Appendix S6 - Table 1: Results from linear regressions with zero-intercepts using validation dataset of observed and predicted summed cover for six plant growth forms and total summed cover transformed using either the predicted mean or Tüxen & Ellenberg (1937) transformations. For each linear regression the fit of the observed and predicted summed cover are assessed using root mean squared deviation (RMSD), slope (with 95% confidence intervals) and adjusted R^2^.

| **Growth form** | **Transformation** | **RMSD** | **Slope** | **Lower slope** | **Upper slope** | **Adjusted R^2^** |
| --- | --- | --- | --- | --- | --- | --- |
| Tree | Tüxen & Ellenberg | 15.49 | 0.91 | 0.91 | 0.92 | 0.98 |
|  | Predicted Mean | 14.97 | 1.08 | 1.07 | 1.09 | 0.97 |
| Shrub | Tüxen & Ellenberg | 6.28 | 0.95 | 0.94 | 0.96 | 0.96 |
|  | Predicted Mean | 5.99 | 1.07 | 1.06 | 1.08 | 0.97 |
| Grass | Tüxen & Ellenberg | 8.58 | 0.86 | 0.85 | 0.87 | 0.94 |
|  | Predicted Mean | 6.13 | 1.07 | 1.06 | 1.08 | 0.96 |
| Forb | Tüxen & Ellenberg | 11.63 | 0.76 | 0.75 | 0.77 | 0.91 |
|  | Predicted Mean | 6.82 | 1.12 | 1.11 | 1.13 | 0.95 |
| Fern | Tüxen & Ellenberg | 5.08 | 0.88 | 0.86 | 0.89 | 0.92 |
|  | Predicted Mean | 4.00 | 1.05 | 1.04 | 1.06 | 0.94 |
| Other | Tüxen & Ellenberg | 2.57 | 0.80 | 0.78 | 0.83 | 0.86 |
|  | Predicted Mean | 1.59 | 1.00 | 0.98 | 1.02 | 0.93 |
| Total | Tüxen & Ellenberg | 27.01 | 0.86 | 0.86 | 0.87 | 0.98 |
|  | Predicted Mean | 20.54 | 1.10 | 1.09 | 1.11 | 0.98 |

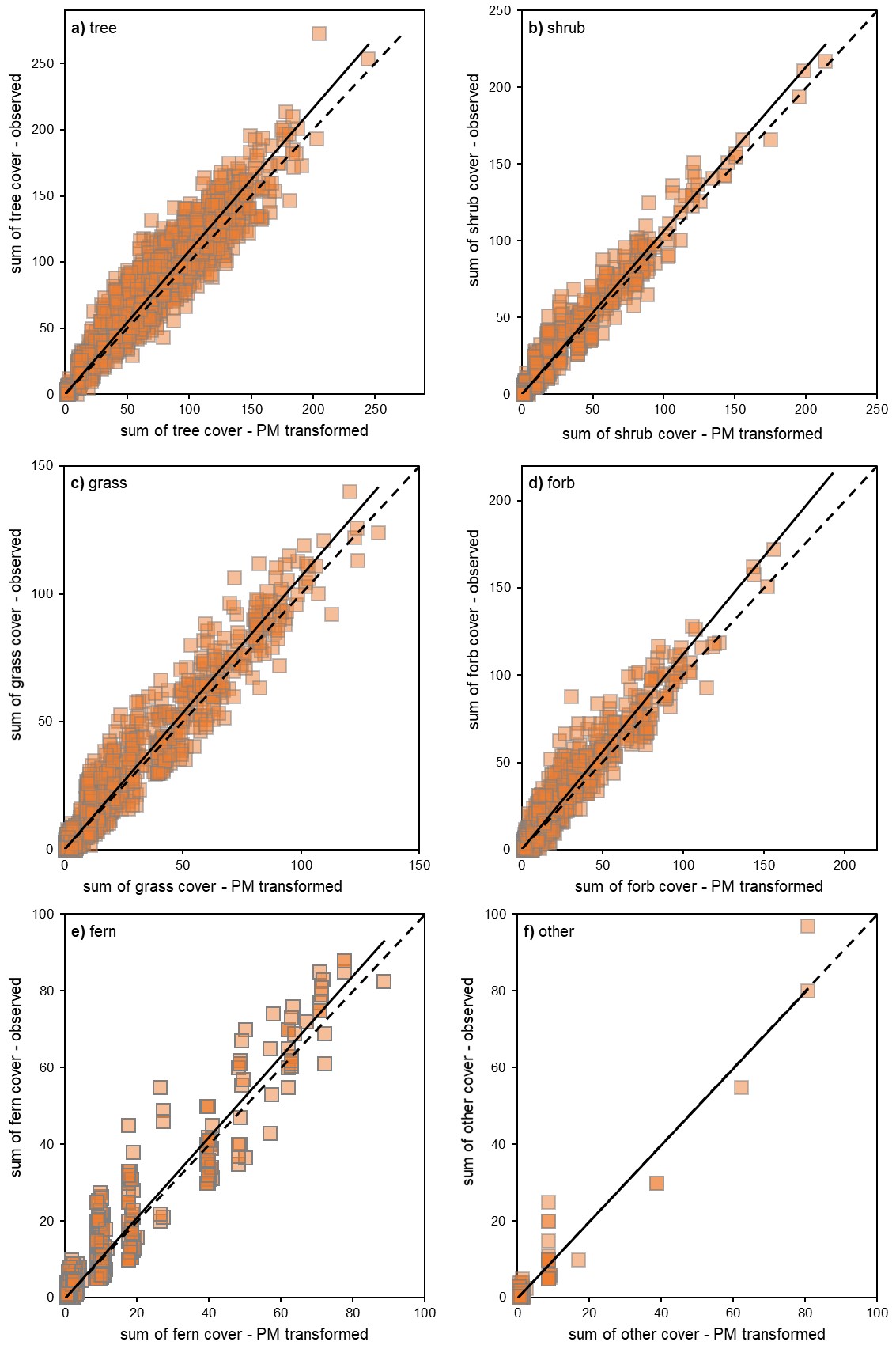

Appendix S7 Figure 1: Scatter plots of validation data showing the relationships between summed cover for the visual estimates of cover (0.01%–100%) compared to the sum of cover when transformed by predicted mean (PM) for different growth forms. Number of observations (n) for **a)** trees (n = 13 850); **b)** shrubs (n = 4 796); **c)** grasses (n = 8 546); **d)** forbs (n = 20 911); **e)** ferns (n = 2 561) and **f)** other (n = 833). The ideal 1:1 line of best fit is shown as dashed line and solid black line shows slope of the regression through the origin. Transformations that overestimate will have a slope <1.

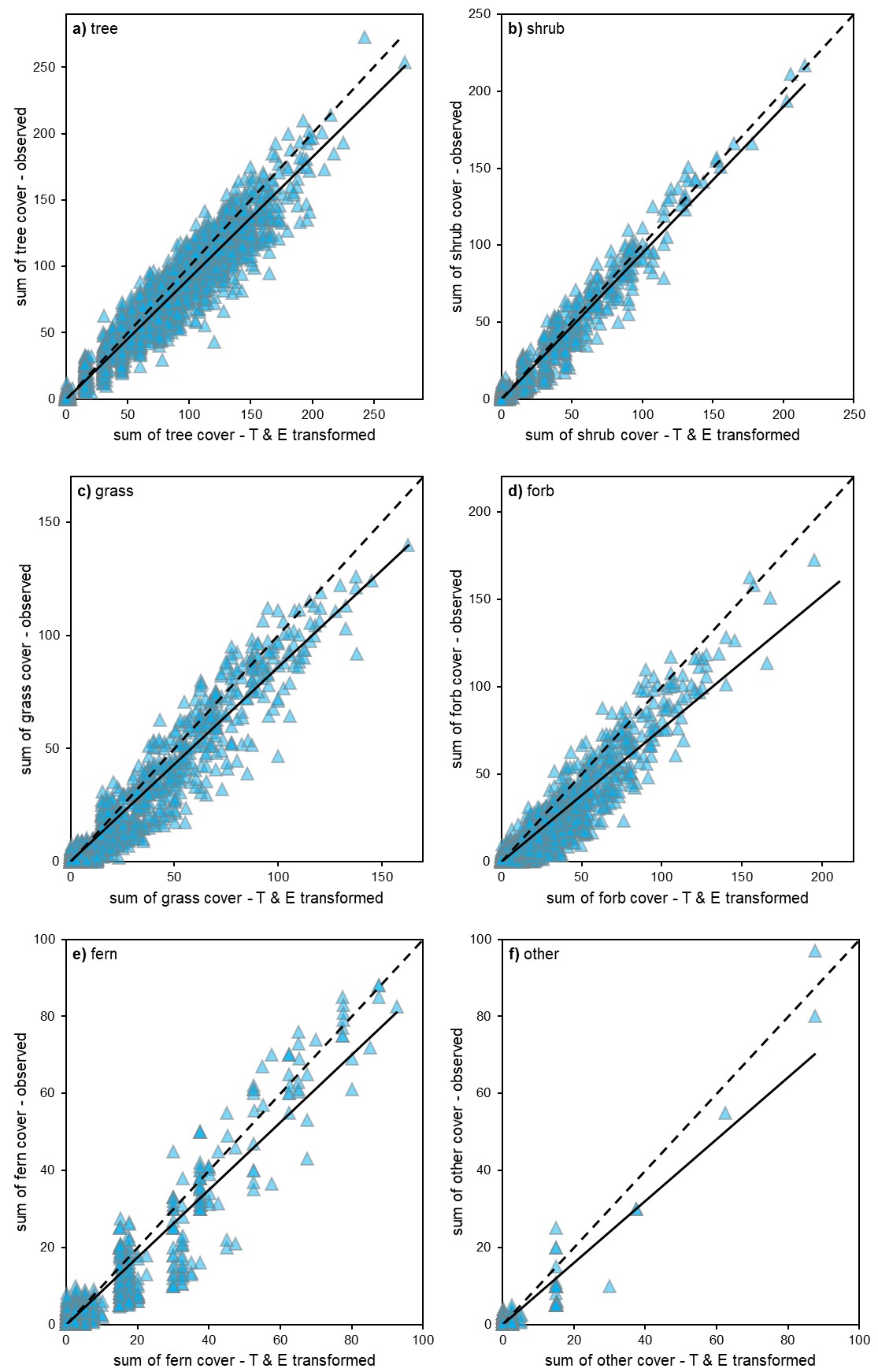

Appendix S7 Figure 2: Scatter plots of validation data showing the relationships between summed cover for the visual estimates of cover (0.01%–100%) compared to the sum of cover when transformed by Tüxen and Ellenberg (1937) (T & E) for different growth forms. Number of observations (n) for **a)** trees (n = 13 850); **b)** shrubs (n = 4 796); **c)** grasses (n = 8 546); **d)** forbs (n = 20 911); **e)** ferns (n = 2 561) and **f)** other (n = 833). The ideal 1:1 line of best fit is shown as dashed line and solid black line shows slope of the regression through the origin. Transformations that overestimate will have a slope <1.
